# Stress fibers are physically and genetically linked to SPIN90 and Arp2/3 complex

**DOI:** 10.64898/2026.07.06.736769

**Authors:** Luther W. Pollard, Cara L. Barnett, Ellie L. Sams, Andrew J. Steen, Jordan R. Beach, Patrick W. Oakes, Qing Tang

## Abstract

The Arp2/3 complex has long been considered to assemble only branched actin structures in the cell (lamellipodia, endocytic patches, comet tails, and many more). We show for the first time by single-molecule tracking (SMT) that the Arp2/3 complex and SPIN90, which activates Arp2/3 complex to nucleate unbranched filaments, bind to and move in the basal cortex with stress fibers (SF) and focal adhesions (FA) that, unlike known sites of Arp2/3 enrichment, employ linear actin bundles. SPIN90 knockout in U2OS cells disrupts the basocortical recruitment of specific Arp2/3 isocomplexes while significantly increasing the speed of collective cell migration, total basocortical Arp2/3 complex retention, nascent FA number, and traction force. Polypeptides containing SPIN90’s SH3 domain show enrichment in FAs and decrease their average size upon overexpression. Thus, our findings show that SF-FA bundles not only directly bind to SPIN90 and Arp2/3 complex; they are also sensitive to genetic manipulations of SPIN90.

**Significance statement:** The Arp2/3 complex, the branched actin nucleator, drives pushing forces in the cytoplasm, yet its diffuse cortical localization has obscured its functions within unbranched structures. In conventional microscopy, associations of molecules with specific structures are impossible to verify when there is little enrichment over the cytoplasm. Whereas other methods fail to provide convincing evidence of such associations, we used single molecule tracking in live cells to identify molecules that track the motions of stress fibers. Thus, we were able to discover that the Arp2/3 complex and SPIN90, proteins whose roles in cell biology have been thus far limited to branched actin networks, directly integrate within the unbranched stress fiber bundles, while genetics support placement of SPIN90 in this pathway.

## Introduction

The Arp2/3 complex, the first actin nucleator discovered, is ubiquitous across Eukarya and is critical for an ever-expanding list of essential cellular functions including leading edge protrusion, phagocytosis, endocytosis, autophagy, organelle trafficking, chromatin repair and remodeling, and meiotic spindle positioning (1–6). The complex consists of seven subunits, <u>a</u>ctin-related <u>p</u>roteins Arp2 and Arp3 as well as <u>Arp</u> <u>c</u>omplex subunits ArpC1-5, which together nucleate filamentous (F)-actin (daughter) from the side preexisting (mother) F-actin (7–12). The Arp2/3 complex activation occurs through Wiskott–Aldrich syndrome protein (WASP)-family nucleation promoting factors (NPF) that prime the complex for daughter filament assembly (13–17). Recently, human SPIN90 (SH3 Protein Interactor of Nck, 90 kDa) and members of its conserved family were shown to bind and activate the Arp2/3 complex at the mother filament-binding interface, resulting in unbranched F-actin (8, 18–23). All members of the SPIN90 family retain a leucine-rich armadillo (ARM) repeat domain that performs this function. Since its discovery, the role of this unbranched form of Arp2/3-actin in mammalian cells has been a mystery.

Because dendritic actin assembly requires preexisting mother filaments off which initial branches can form, one model is that SPIN90 family proteins help to initialize dendritic actin assembly through mother filament nucleation, and SPIN90 has been shown to promote dendritic assembly in lamellipodia in certain contexts (24–28). However, a pervasive network of F-actin in the cortices of many animal cells challenges the necessity for mother filaments to be generated in such a manner. SPIN90 amongst Holozoa contains an SH3 domain (that binds proline rich sequences) and a dimerization domain (DD) that do not exist in fungal homologues (21, 22), indicating differences in the molecular and cellular functions amongst this family. In contrast to the *S. pombe* yeast homolog Dip1 (24), SPIN90 is not only enriched in dendritic structures; it also localizes more diffusely in the cortex of mammalian cells where it has implicated functions in cell shape, cortical stiffness, cell-substrate adhesion (26–30). Since it is established that SPIN90-Arp2/3 complexes assemble unbranched F-actin, there remains the possibility that the Arp2/3 complex also can function in linear actin structures.

The Arp2/3 complex is a major component of the cortical actin network that regulates its density and mechanics (31, 32). Ventral stress fibers (SF) are unbranched bundles in the cortical actin network at the base of the cell that, together with integrin-mediated anchoring by focal adhesions (FA), provide traction between the cell and the substrate (33). SF-FA bundles generate tension with contractile units formed from alternating nonmuscle myosin-II and F-actin crosslinking protein α-actinin (34–36). Historically, the Arp2/3 complex has not been considered part of SF-FA bundles. Rather, it has been implicated to siphon actin away from SFs and into competing branched networks (37). However, SFs that are formed by myosin-II’s coalescence of lamella-derived filaments, or transverse arcs, are only assembled when the Arp2/3 complex is expressed (36). Whether transverse arcs contain Arp2/3-nucleated actin or are simply a byproduct of the cellular geometry induced by lamellipodia remains to be shown. In general, it has been unclear where the distinction lies between SFs and the cortex at large, and if Arp2/3 complex is truly “competitive” with SFs, then what are the mechanisms to exclude it from entering SFs?

To gain new insight into the biology of the Arp2/3 complex, SPIN90, and their associated unbranched actin filaments in the cell cortex, here we employ live-cell single molecule tracking (SMT) super-resolution microscopy. Our findings reveal that the Arp2/3 complex and SPIN90 enter into and regulate SF-FA bundles. SPIN90 ablation leads to increased cortical Arp2/3 complex recruitment, increased cellular tension, and an increased number of smaller adhesions highlighting critical roles for SPIN90-Arp2/3 complexes within linear bundles.

## Results

While a body of evidence points to the Arp2/3 complex regulating the assembly of SFs and FAs, direct evidence of Arp2/3 complex incorporation into unbranched structures in the cell is lacking (36, 38, 39). While we sometimes see an apparent enrichment of Arp2 overlapping with SF-dense regions by immunofluorescence (IF; Fig. 1A), we sought parallel evidence to support our observation. Therefore, we asked whether the Arp2/3 complex could be directly visualized in SF-FA bundles. We reasoned that SMT should provide robust evidence of structural association by showing molecules entering the bundles and tracking their motion. The advantage of SMT over other approaches is that the confined, diffusive, or directed nature of trajectories offers an enhanced view of the biochemical activities of individual molecules localized with high-precision in living cells. If a protein maintains a population localized broadly within the cell cortex but can be subdivided into active/bound versus inactive/unbound pools, this SMT approach can readily distinguish the two. However, the SMT approach is poor at quantifying the concentrations or relative densities of the target proteins.

**Fig. 1.**
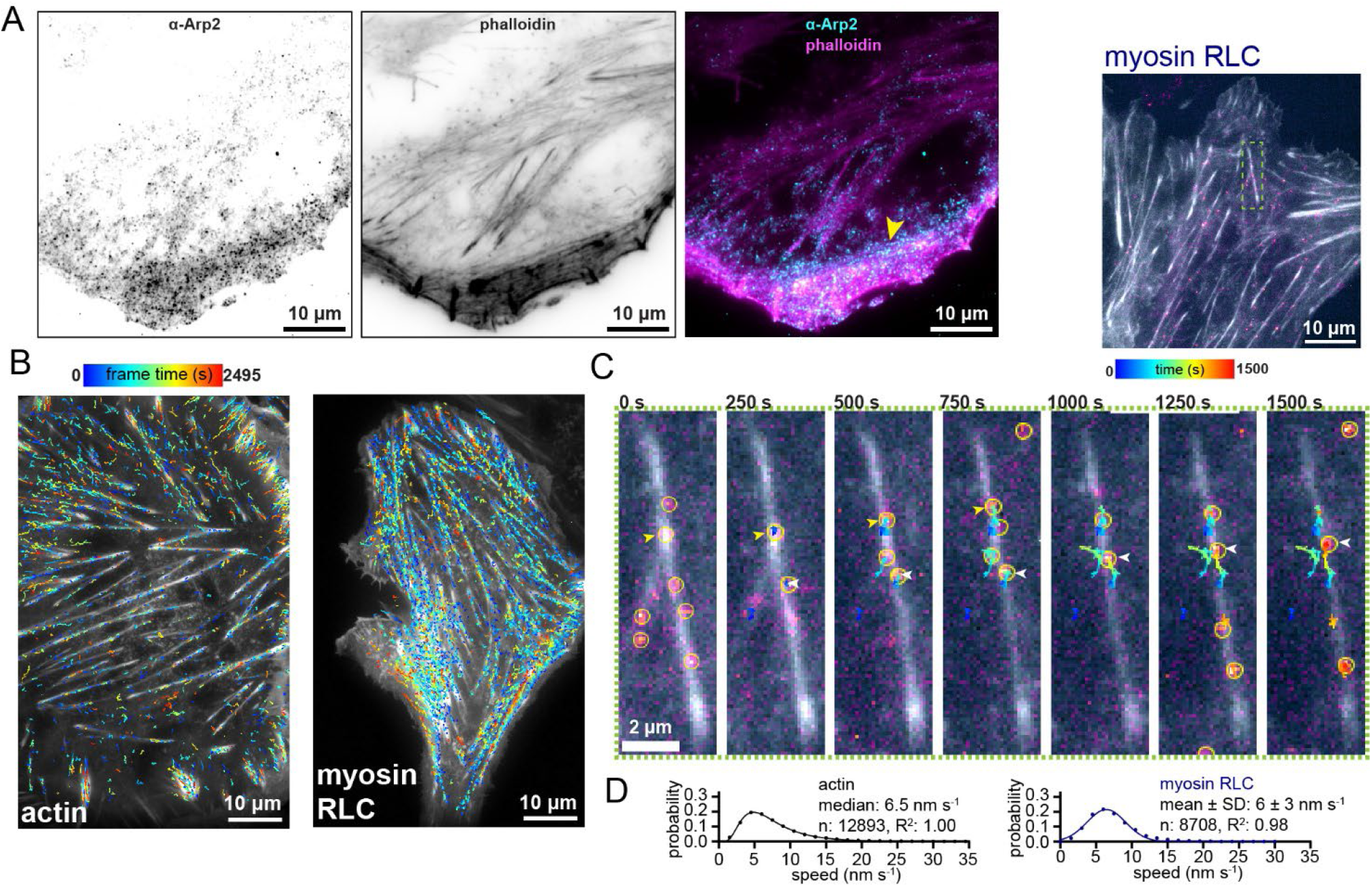
Live cell single molecule tracking (SMT) of actin and myosin-II in stress fiber-focal adhesion (SF-FA) bundles. (**A**) U2OS fixed, permeabilized, and immunostained for Arp2 and actin (phalloidin). Example shows a cell where SF density correlates with Arp2 signal, particularly in transverse arcs (arrowhead). (**B-D**) U2OS cells ectopically expressing LifeAct-EGFP and Halo fusion proteins stained at single molecule level with JaneliaFluor-JFX^650^ Halo ligand. Molecular trajectories are from image series acquired every 5 s using total internal refection fluorescence (TIRF) microscopy (*Materials and Methods*). (**B**) Representative maximum projections of GFP fluorescence over 500 frames overlaid with trajectories of the indicated Halo fusion proteins lasting at least 2 min. (**C**) Example cell (top right) and time series (inset, below) show Halo-labeled single molecules and their associated trajectories moving in stress fibers (colored arrowheads point to different molecules). Localizations (yellow circles), LifeAct-EGFP (grayscale), and Halo signal (magenta to yellow intensity scale) are shown. (**D**) Speeds of molecules with directed motion (MSD α ≥ 1.2). (Symbols) frequency and (solid lines) lognormal (actin) or normal (RLC) fits. Median or mean ± SD speeds are reported depending on distribution. Number of biological repeats (N_BR_; experiments performed on different days): (actin) 9 and (RLC) 5. Number of trajectories (n) are shown.

To identify proteins by SMT in cortical SF-FA bundles we used human osteosarcoma U2OS, an established cell line for studying these structures (36, 40–42), and expressed Halo-tagged proteins that are known components SF-FA bundles, including actin, myosin-II regulatory light chain (RLC), α-actinin (36), and Nck1 (an SH2-SH3-containing FA adapter protein) (43). These constructs were expressed exogenously alongside LifeAct-EGFP and stained with sub-saturating (1 nM) Janelia Fluor-JFX^650^ Halo ligand (to achieve single-molecule labeling, see *Materials and Methods*) and then imaged in live cells using total internal reflection fluorescence (TIRF) microscopy. Images were acquired at 0.2 Hz with a threshold for trajectory analysis of ≥ 2 min duration in order to capture single molecules associated with SF traction, which typically have speeds < 30 nm s^-1^ (44), and to filter out faster dynamics such as lamellipodial turnover (45, 46). Trajectories were then subjected to motion classification (directed, diffusive, and confined) by mean squared displacement (MSD) analysis (*Materials and Methods*). Over several minutes, the trajectories of these proteins overlapped and tracked with SF-FA bundles (Figs. 1B and C, S1A-D). On this timescale, SFs were highly dynamic (coalescing and disassembling) and had variable thicknesses and intensities, which precluded us from making a reliable SF reference mask to colocalize trajectories of Halo-proteins. Thus, we analyzed trajectories of the Halo-proteins globally within the basal cortex, ∼100 nm deep by TIRF. Particles with directed motion (MSD α ≥ 1.2) moved with a broad distribution of speeds associated with assembly and contraction of bundles (Figs. 1D, S1B and D), approximately matching the lognormal speed distribution of actin traction obtained by Gardel *et al.* (44).

CAPZ, hereafter referred to as “capping protein”, caps the fast-growing barbed end of F-actin, controlling its net polymerization in the cytoplasm (47). Investigators were previously able to show through speckle microscopy that capping protein enters the SFs (48), even though its localization in the cell body is typically diffuse. By our SMT method, capping protein exhibits a mixed population of trajectories, some of which track together with the bundles while others are elsewhere in the cortex (Fig. S1G and H). In contrast, Halo by itself did not show specific bundle tracking but had sporadic trajectories near the cortex (Fig. 2A and D; Movie S1). Directed capping protein trajectories had a normal speed distribution with an average of 6 ± 3 nm s^-1^ (standard deviation, SD), the same as RLC, α-actinin, and Nck1 (Figs. 1D and S1A-H). Thus, our SMT approach corroborates the previous study’s findings (48).

**Fig. 2.**
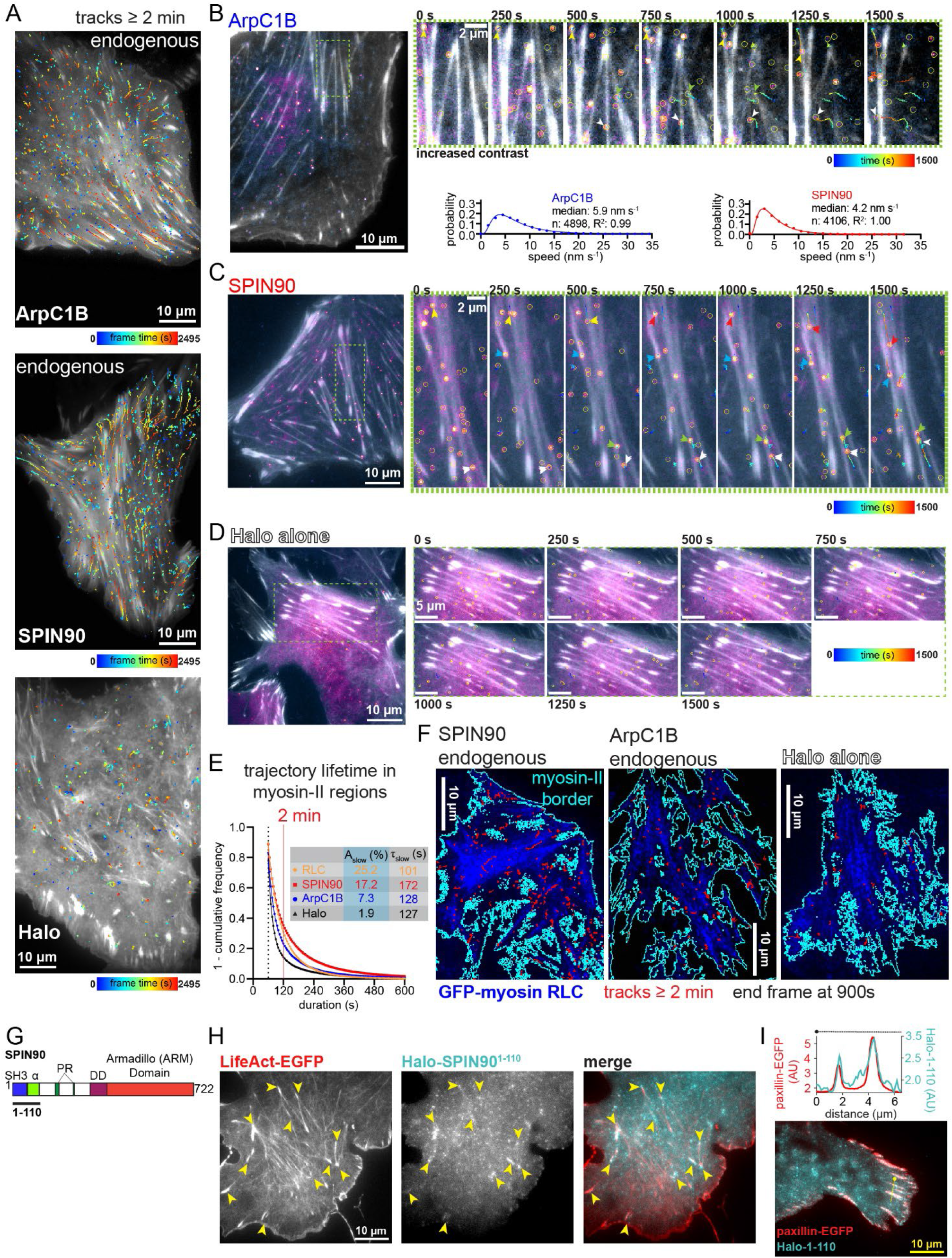
Live cell SMT of Arp2/3 complex, SPIN90, and Halo alone. Using SMT methods employed in Fig. 1, Halo-proteins were tracked in SF-FA bundles. (**A**) Maximum projections of LifeAct-EGFP signal over 500 frames overlaid with trajectories of the indicated Halo fusion proteins lasting at least 2 min. Representatives for Halo-ArpC1B and Halo-SPIN90 are tagged at their native loci (*Materials and Methods*). (**B-D**) Example cells (left) and time series (insets, right) showing Halo-labeled molecules and their associated trajectories moving in stress fibers (arrowhead color indicates individual molecules). Representatives show ectopically expressed Halo fusions: (**B**) ArpC1B, (**C**) SPIN90, (**D**) Halo alone. Localizations (yellow circles), LifeAct-EGFP (grayscale), and Halo signal (magenta to yellow intensity scale) are shown. For better visualization of thin F-actin fibers, contrast is increased for the inset as indicated. Speeds of molecules represented as in Fig. 1D (below). Data are from N_BR =_ (ArpC1B) 13 (10 with ectopic expression and 3 with endogenous Halo-tagging) and (SPIN90) 16 (13 ectopic and 3 endogenous), with (n) number of trajectories. (**E**) Survival of trajectories (tracks) overlapping with clusters of myosin RLC (GFP-RLC or Halo-RLC by itself) plotted as 1 – cumulative frequency (points) and fit to double exponentials (solid lines). The amplitudes (A_slow_) and lifetimes (τ_slow_) are shown. A cutoff of 12 frames (dashed line) was employed to reduce noise from diffusive molecules. Unfused Halo is distinct from the other conditions at an inflection around 2 min (red line). Statistics in Table 1. (**F**) Examples taken from data in (**E**) where trajectories (red) lasting at least 2 min are overlaid on top of the final frame (900 s) of the GFP-RLC channel (blue), showing the relative density of long-lived trajectories. (**G**) SPIN90 domain diagram. (**H** and **I**) Panels show representative cells where (**H**) Halo-SPIN90^1-110^ has FA-like enrichments at the ends of SFs (yellow arrowheads) or (**I**) colocalize with paxillin-EGFP (maximum projection image: 90 s of continuous acquisition at 2 Hz). Line-scan intensity plot of Halo-SPIN90^1-110^ and paxillin-EGFP shown.

**Table 1.** Lifetime (τ) kinetics of Halo-protein trajectories in myosin-II regions.

|  | best fit values (95% confidence interval) |  |  | N<sub>BR</sub> |  |  |  |
|---|---|---|---|---|---|---|---|
|  | τ<sub>slow</sub> (s) | A<sub>slow</sub> (%) | τ<sub>fast</sub> (s) | total (endogenous) | cells | tracks | R<sup>2</sup> |
| **myosin-II RLC** | 101 (99-103) | 25.2 (24.5-25.9) | 34.6 (33.8-35.4) | 7 | 14 | 96,112 | 1.0000 |
| **SPIN90** | 172 (167-177) | 17.2 (16.8-17.6) | 32.7 (31.6-33.6) | 7 (3) | 39 | 30,242 | 0.9998 |
| **ArpC1B** | 128 (124-132) | 7.3 (7.1-7.6) | 23.0 (22.2-23.7) | 5 (3) | 34 | 15,754 | 0.9997 |
| **Halo alone** | 127 (122-134) | 1.9 (1.8-2.0) | 17.4 (16.9-17.9) | 4 | 24 | 9,189 | 0.9995 |

Halo fusion proteins of Arp2/3 complex (Halo-ArpC1B) as well as its effectors SPIN90 and cortactin, a protein that binds and stabilizes branched and unbranched Arp2/3-actin at the daughter filament junction (23, 49, 50), were frequently seen to bind and move together with SF-FA bundles in SMT experiments (Figs. 2A-C, S1I and J; Movies S2-5). Both Arp2/3 complex and SPIN90 had SF-FA bundle binding and tracking behaviors whether exogenously expressed or endogenously tagged at their native loci in U2OS (*ARPC1B* for the Arp2/3 complex; Figs. 2A-C, and S2; Movies S2-5). These behaviors were also seen upon ectopic expression of Halo-tagged ArpC1B and SPIN90 in different cell lines: mouse fibroblast-derived cells (MEF and NIH-3T3) and human cervical cancer HeLa cells (Movies S6 and S7). N-terminally tagged ArpC1B was previously shown to assemble complete complexes, localize normally in cells, and have full functionality *in vitro* (48, 51, 52). Accordingly, Halo-ArpC1B is enriched and undergoes retrograde flow in the lamellipodial leading edge (Fig. S3), indicating normal Arp2/3 complex assembly. To determine kinetic differences in SF binding between Halo-fusions of myosin-II RLC, ArpC1B, and SPIN90 compared to Halo alone, we co-expressed them with GFP-RLC (except for Halo-RLC) and analyzed trajectory lifetimes within RLC-dense regions. The resulting lifetime frequency distributions best fit two-phase decay models (Fig. 2E, Table 1). Based on these fits, the amplitudes (A_slow_) of the slow dissociation rates (k_slow_) of Halo-tagged RLC, SPIN90, and ArpC1B were 4-13-fold greater than Halo alone (Fig. 2E, Table 1), suggesting that these molecules have more long-lasting trajectories while the unfused Halo population is dominated by transient associations. Consequently, we found that using a cutoff of 120 s or above for trajectory lifetime could robustly differentiate the cortical binding of different Halo-proteins (Fig. 2E and F). SPIN90 associated more strongly within myosin-II-dense regions than ArpC1B by the average trajectory lifetime (172 versus 128 s), A_slow_ (17.2 versus 7.3%), and number of trajectories above the cutoff (Fig. 2E and F, Table 1). SPIN90 and ArpC1B molecules also differ slightly in the speed of their directed motions (4.2 versus 5.9 nm s^-1^; Fig. 2B and C). It is possible that the lifetime and speed discrepancies between SPIN90 and ArpC1B come from SPIN90 binding the cortex independently of the Arp2/3 complex (through different domains), which we next examined.

We next dissected how SPIN90’s domains differentially contribute to basocortical binding by expressing various Halo-tagged segments of SPIN90 in SPIN90 knockout (KO) cells (*Materials and Methods*; Fig. S4) with or without an ARM domain mutation (ARM*), E588K R645E, that abolishes Arp2/3 complex activation *in vitro* (19). We anticipated that SPIN90 mutants that are defective in recruitment to the basal cortex or Arp2/3 complex binding would result in reduced association of SPIN90 to the basal cortex. However, SPIN90 mutants re-expressed in SPIN90 KO cell lines had variable protein levels from cell to cell, despite having comparable expression levels at the population level (Fig. S4), preventing relative affinity determination by signal intensity alone. Hence, we made the approximation that long-lasting SMT trajectories are proportional to the amount of stable binding and that localization number is proportional to concentration. The relationship between these parameters gave linear correlations that varied by protein, the relative slopes of which were used to extract a basal cortical binding index (BCBI; *Supplemental Materials and Methods*). Among the fragments tested, Halo-SPIN90^1-110^, containing the SH3 domain, showed the highest basocortical binding and was enriched in FAs (Figs. 2G-I, S5). Halo-SPIN90^269-722^, which contains the DD-ARM domains and activates actin nucleation (19, 21, 22), also was found to bind robustly the basal cortex except when the DD domain was deleted or it contained the ARM* mutation (Fig. S5A and D). These results suggest that SPIN90 has at least two mechanisms of basocortical association: through its SH3 and through Arp2/3 complex activation.

The discovery of the Arp2/3 complex and SPIN90 in SF-FA bundles invoked the question of whether SPIN90-KO cells exhibit defects in cell migration, traction, and substrate adhesion. In wound-healing assays, two SPIN90-KO clones independently had increased migration speeds, 19 ± 1 µm h^-1^ and 18 ± 4 µm h^-1^, compared to the speed of their unedited isogenic U2OS cell line (WT), 9 ± 1 µm h^-1^ (*p* = 0.0143, Fig. 3A-C). We hypothesized that this effect could be related to actomyosin speed and traction stress (44). SMT analysis of Halo-actin showed that both SPIN90-KO cell lines had reduced geometric mean and maximum speeds compared to WT (Fig. 3D-F). Halo-RLC speeds in the SPIN90-KO lines were also reduced more than 20% from the WT speed (Fig. 3G-I and K). Moreover, the ratio of directed to confined (MSD α ≤ 0.8) trajectories in each of the SPIN90-KO lines was less than 20% of the WT ratio (Fig. 3J and K), demonstrating myosin particles in KO cells were more static in addition to having slower directed motion. To examine how these changes were related to the cell’s ability to generate force, we performed Traction Force Microscopy (TFM). TFM experiments revealed that both SPIN90-KO lines exert more than double the contractile energy per area than the WT (Fig. 3L-N), despite remaining approximately the same size (Fig. 3N). The reduction in SF particle motion in SPIN90-KO cells is therefore best explained by the inverse relationship of force and velocity for contractile myosins (53, 54). Re-expression of Halo-SPIN90, however, failed to restore traction force in the KO lines down to WT levels (Fig. 3L-N), suggesting that SFs in the KO lines have adapted long-term to the absence of SPIN90. Nonetheless, these data highlight the importance of the SPIN90 gene to SF-FA mechanochemistry.

**Fig. 3.**
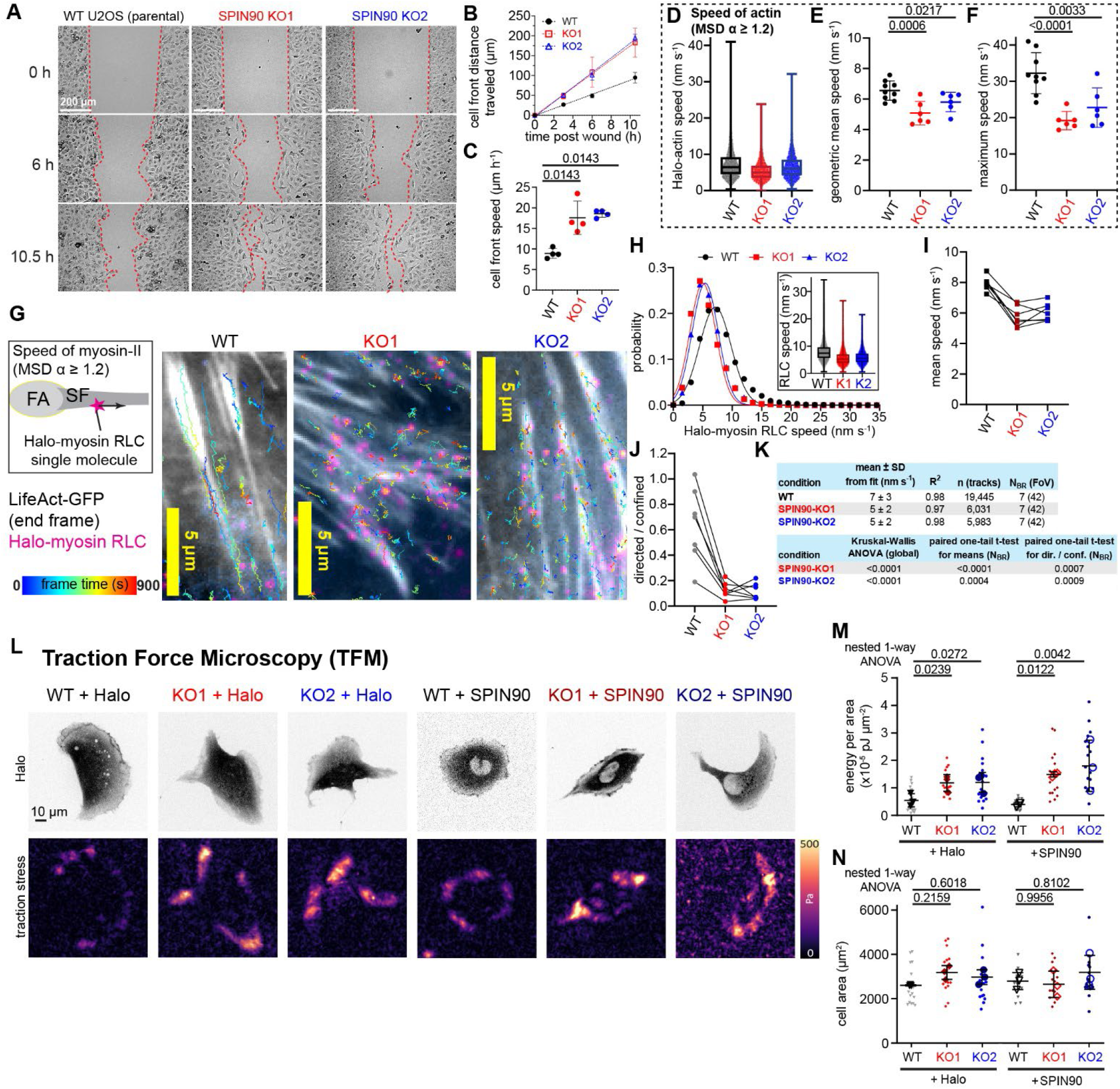
Effects of SPIN90 ablation on cell motility and directed actomyosin motion in U2OS. (**A-C**) Two SPIN90-KO clones and their unedited U2OS parent cell line (WT) were assessed for their ability to migrate across a gap produced by removal of a barrier in the culture dish. Data are from N_BR_ = 4 experiments. (**A**) Example widefield micrographs showing gap closure over time. Yellow dashed lines trace the edges of the wound. (**B**) Cell front distance traveled over time averaged globally. Shown are means (symbols), SD (bars), and linear fits (dashed lines) with slopes (± standard error, SE) of (WT) 9 ± 1 µm h^-1^, (KO1) 18 ± 2 µm h^-1^, and (KO2) 19 ± 1 µm h^-1^. (**C**) Cell front speed. Slopes from each biological repeat (symbols) and the overall mean ± SD (lines and error) are shown. *P* values are calculated by one-tailed Mann-Whitney test. (**D-F**) SMT speeds of Halo-actin molecules with directed motion in WT and SPIN90-KO U2OS. (**D**) Global speed distributions. The speeds of individual trajectories (symbols; WT, n = 12893; KO1, n = 7205; KO2, n = 7557) as well as the median, interquartile range, and min-max values (box and whiskers) are shown. The frequency distribution of the WT data is shown in Fig. 2B. (**E**) Geometric mean and (**F**) maximum speeds. (**E** and **F**) Values from each biological repeat (symbols) and the overall mean ± SD (lines and error) are shown. *P* values are calculated by unpaired, one-tailed t-test. (**G-K**) Speeds of directed Halo-myosin RLC molecules by SMT. (**G**) Cartoon diagram and example trajectories. (**H**) Frequency distributions with (solid lines) Gaussian fits and (inset) box (interquartile range) and whiskers (min. and max.) plots showing all points (speeds of individual trajectories). (**I**) Mean speeds (points) connected by solid lines showing each experiment. (**J**) The ratio of directed (MSD α ≥ 1.2) to confined (α ≤ 0.8) trajectories of each experiment (points and lines). (**K**) Statistics (*p* values) for **G-J**, number of fields of view (FoV) shown. (**L-N**) Traction force microscopy (TFM) experiments comparing WT to SPIN90-KO lines. (**L**) Representative images of U2OS expressing Halo alone or Halo-SPIN90 and corresponding traction stress maps. Plots of (**M**) contractile energy per area and (**N**) cell area show (small symbols) individual cells, (large symbols) replicate means, (horizontal lines and error bars) overall mean ± SD of the N = 3 replicate means. (**M** and **N**) TFM data *p*-values: nested one-way ANOVA of log₁₀-transformed data followed by Tukey’s multiple comparisons test.

To examine the effects of SPIN90 ablation on FAs, we performed TIRF-IF microscopy with fixed WT and SPIN90-KO U2OS cells staining for proteins that assemble nascent FAs, paxillin and phosphorylated focal adhesion kinase (pY397-FAK), as well as proteins that are recruited only to mature FAs, phosphorylated vinculin (pY100) and zyxin (Figs. 4 and S6). Paxillin staining intensity in FAs was reduced in the SPIN90-KO clones by more than 20% (Fig. 4A and B). There were additionally small, but significant differences in FA area and number: the mean area of individual paxillin FAs in SPIN90-KO cells was reduced by 0.1 µm^2^ compared to WT (Fig. 4C), corresponding to SPIN90-KO cells having 1 more FA every 10 µm^2^ total paxillin area (Fig. 4D), together suggesting that the total FA area is divided into more numerous, smaller adhesions consistent with an increase in nascent FAs. This increase was also shown by SPIN90 ablation resulting in average reductions in both pY100-vinculin (30%) and pY397-FAK (20%) FA staining (Fig. S6). An increase in nascent FAs is consistent with the KO lines being more migratory than the WT (Fig. 3A-C). We found that expression of Halo-SPIN90^FL^ in SPIN90-KO cells, but not Halo-SPIN90^1-305^ or HaloTag alone, increased the mean size of zyxin-stained FAs into a regime more similar to that of WT cells expressing Halo alone (Fig. S4E-H). Thus, we observe that SPIN90 re-expression suppresses the production of nascent FAs.

**Fig. 4.**
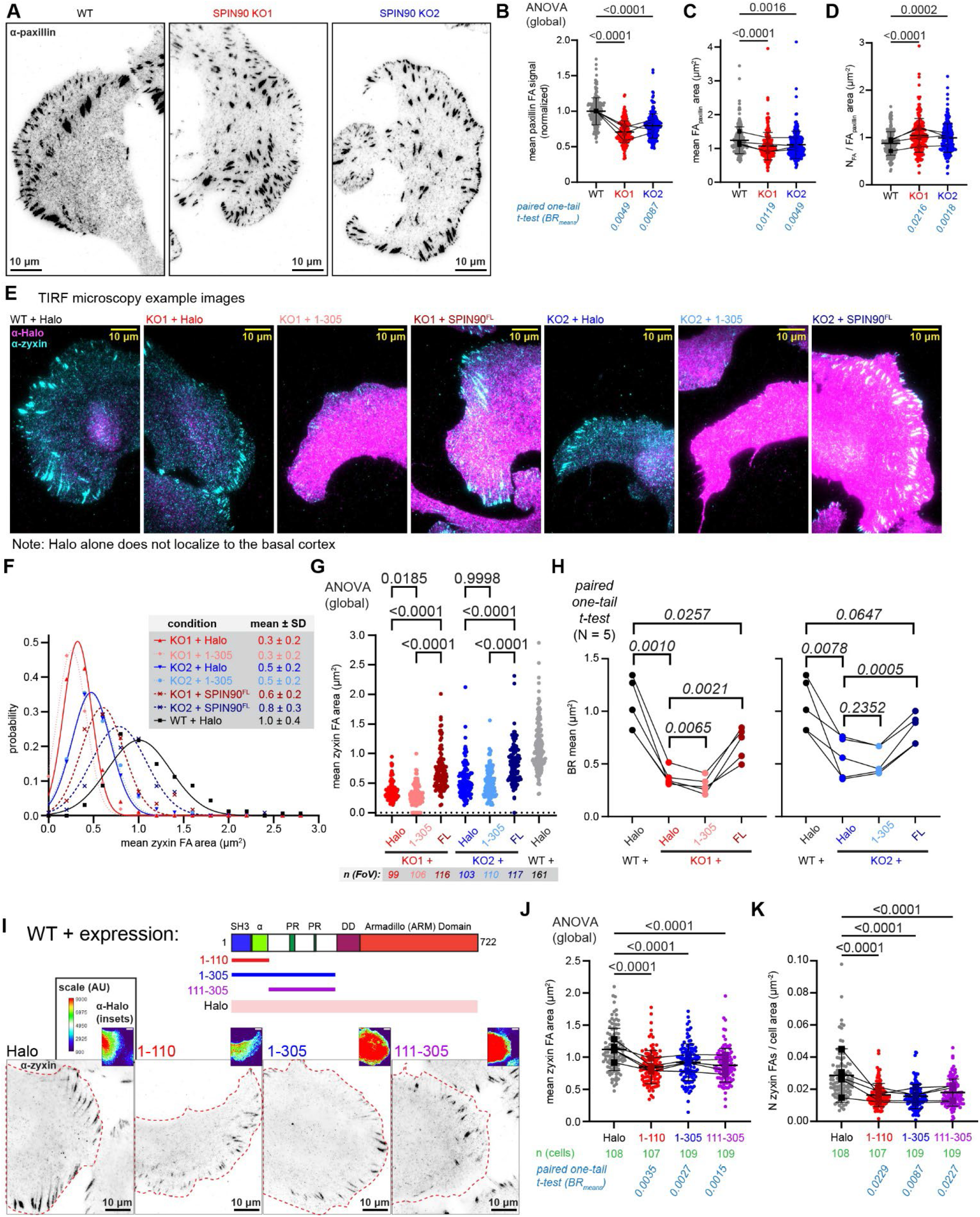
Effects of SPIN90 ablation and expression on FA size. WT and SPIN90-KO U2OS were fixed, permeabilized, immunostained for paxillin, zyxin, or HaloTag, and subjected to TIRF microscopy. (**A**) Example cells stained for paxillin (Alexa^488^). FAs were analyzed for (**B**) paxillin intensity, (**C**) FA area, (**D**) number of FAs (N_FA_) per total FA area, and Data (**B-D**) are from N_BR_ = 4 experiments with n = 202-206 images per condition. (**E-H**) WT or SPIN90-KO U2OS expressing Halo alone, Halo-SPIN90^1-305^, or Halo-SPIN90^FL^ subjected to IF analysis. (**E**) Example cells imaged in TIRF showing (magenta, Alexa^647^) HaloTag and (cyan, Alexa^488^) zyxin staining. (**F**-**H**) Analysis of the mean FA size from zyxin signal in each field of view (FoV). (**F**) Frequency distributions with normal fits (lines) yielding means and SDs. (**G**) Mean FA areas per FoV (points) compared globally by ANOVA (*p* values). (**H**) Means from biological repeat (BR) datasets analyzed by paired one-tail t-test (*p* values). Number of BR (N) and FoV (n) are shown. (**I-K**) WT cells were transduced to express Halo-proteins for 48 h preceding fixation. (**I**) Diagram and example cells stained for zyxin (Alexa^488^) and HaloTag (Alexa^647^; insets with intensity scale, bars: 10 µm). FA maturation was analyzed by (**J**) mean area of FAs stained by zyxin and (**K**) number of zyxin-containing FAs per cell area. (**J** and **K**) Data are from N_BR_ = 5 experiments with (n) number of cells shown. (Graphs) Mean values for each image (circles), means of each biological repeat (BR_mean_; squares), and the global mean ± SD (horizontal lines and error bars) are shown. BR_mean_ values are connected to show each experiment (solid lines). (Statistics) *P* values are calculated by one-way ANOVA, comparing global distributions, and by pairwise one-tailed t-tests.

To probe SPIN90’s phenotypic interactions with FAs more acutely, we ectopically expressed Halo-tagged SPIN90 N-terminal segments in WT U2OS and performed IF-TIRF microscopy with α-zyxin and α-Halo antibodies (Fig. 4I). The mean area of zyxin FAs was reduced in cells expressing SPIN90^1-110^ (difference: -0.27 ± 0.04 µm^2^), SPIN90^1-305^ (-0.20 ± 0.04 µm^2^), and SPIN90^111-305^ (-0.25 ± 0.04 µm^2^) compared to cells expressing unfused Halo (mean: 1.1 ± 0.3 µm^2^; Fig. 4J). The number of zyxin positive FAs per cell area in cells expressing N-SPIN90 fragments was 49-55% of cells expressing Halo alone (Fig. 4K). These findings show that the size and number of mature FAs can be directly linked to SPIN90 activity.

We questioned whether SPIN90’s activation of the Arp2/3 complex in the basal cortex could potentially bridge SPIN90’s effects on traction and adhesion. We performed IF on WT and SPIN90-KO cells staining Arp2, an obligate subunit, and F-actin with AlexaFluor^647^-phalloidin. Arp2 basocortical staining of SPIN90-KO lines was more than double that of the WT (Fig. 5A and B), indicating that total Arp2/3 complex recruitment to the basal cortex is increased by SPIN90 ablation. SPIN90-KO also resulted in a 20 or 50% increase in basocortical phalloidin stain (Fig. 5C). A recent report showed that SPIN90 enhances the enrichment of specific ArpC5L-containing isotypes of the Arp2/3 complex at the lamellipodial leading edge (LE) in mouse melanoma cells (28). Thus, we examined whether SPIN90 has a similar function in the basal cortex. Although ArpC5L staining did not seem to have FA-specific enrichment (Fig. 5D), basocortical ArpC5L signal, including regions overlapping with FAs, in SPIN90-KO lines was less than 50% of the WT signal (Fig. 5E). ArpC5L LE signal in SPIN90-KO clones was 50% of the WT (Fig. 5F), consistent with the aforementioned study (28). Western blots of WT and SPIN90-KO cytoplasmic extracts staining ArpC5L, Arp2, and Arp3 showed no difference in complex subunit expression (Fig. 5G), suggesting that it is recruitment, not expression, that is altered. The increase in basocortical Arp2 and phalloidin staining resulting from SPIN90-KO (Fig. 5A-C) suggested that the Arp2/3 complex retention and F-actin assembly in the cortex becomes denser in the absence of SPIN90, as shown in work from Cao *et al.* (29). We therefore sought to determine if the Arp2/3 complex binds SFs in SPIN90-KO cells. Indeed, we found that Halo-ArpC1B retains its ability to bind SFs in the SPIN90-KO (Fig. 5H). Thus, our data show that SPIN90 regulates the density and composition of the Arp2/3 complex in the basal cortex but is not required for its binding to SF-FA bundles.

**Fig. 5.**
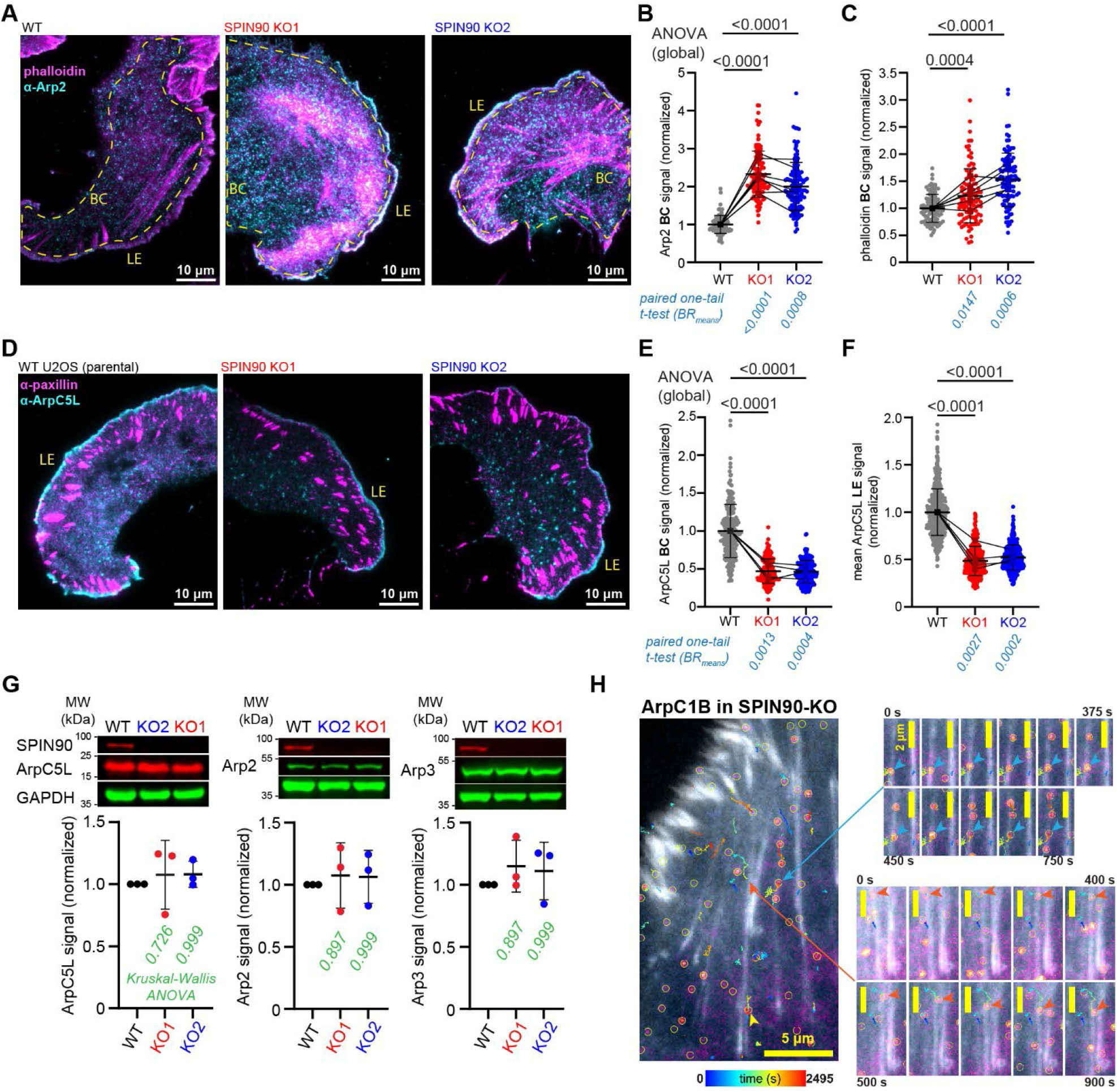
Effects of SPIN90 ablation on Arp2/3 complex isotypes by IF analysis. (**A-F**) WT and SPIN90-KO U2OS were fixed, permeabilized, and stained for F-actin (Alexa^647^-phalloidin), Arp2, paxillin, and ArpC5L. (**A**) Representative TIRF images of cells co-stained for phalloidin (magenta) and Arp2 (cyan, Alexa^488^). Lamellipodial leading edge (LE) and basocortical (BC; dashed lines) regions indicated. The basal cortex was analyzed for (**B**) Arp2 (N_BR_ = 7, n = 120 cells) and (**C**) phalloidin (N_BR_ = 6, n = 100 cells) intensities. (**D**) Representative TIRF images of cells co-stained for paxillin (magenta, Alexa^488^) and ArpC5L (cyan, Alexa^647^). ArpC5L intensities in the (**E**) basal cortex (N_BR_ = 4, n = 202-206 cells) and (**F**) leading edges (N_BR_ = 4, n = 318-493 LEs) were measured. (Graphs) Mean values for each image (circles), means of each biological repeat (BR_mean_; squares), and the global mean ± SD (horizontal lines and error bars) are shown. BR_mean_ values are connected to show each experiment (solid lines). (Statistics) *P* values are calculated by one-way ANOVA, comparing global distributions, and by pairwise one-tailed t-tests. (**G**) WT and SPIN90-KO cytoplasmic extracts are analyzed for SPIN90, ArpC5L, Arp2, and Arp3 levels by immunostaining of Western blots, using GAPDH as a loading control. Data are from N_BR_ = 3 experiments. (Top) Representative Western blots showing WT expression and absence of SPIN90 in KO clones. (Bottom) Band intensities (symbols) normalized to both the WT and GAPDH loading controls, means ± SD (lines and bars) and ANOVA *p* values are shown. (**H**) Example trajectories of Halo-ArpC1B tracking SF movements in a SPIN90-KO cell. Localizations (yellow circles), LifeAct-EGFP (grayscale), and Halo signal (magenta to yellow intensity scale) are shown. Colored arrows/arrowheads point to different molecules bound to fibers. Time series bars: 2 µm.

## Discussion

Here we make the discovery that, in addition to their established role driving dendritic actin assembly, the Arp2/3 complex and SPIN90 incorporate into SF-FA bundles, which has major implications for mechano-architectural models of how these structures are functionally regulated. In the absence of SPIN90, U2OS cells show increased recruitment of Arp2/3 complex to the basal cortex, increased numbers of nascent FAs, and become more migratory while exerting higher traction forces. These phenotypes impacting SF-FA bundles are consistent with SPIN90 regulating these unbranched structures, which we speculate results from a direct physical interaction. Since SPIN90 is known to stimulate Arp2/3 complex to nucleate unbranched actin *in vitro* (8, 18–23), it is plausible that such filaments could mediate this regulation.

New high-resolution structures show that homodimerization of human SPIN90 through a high-affinity site is essential for SPIN90’s NPF mechanism, in which each Arp2/3 complex makes critical contacts with both SPIN90 chains (21, 22). This organizational feature of SPIN90 has interesting potential to augment the geometry of actin in SF-FA bundles. While SPIN90 recruits ArpC5L-containing Arp2/3 complexes to the cortex and leading edge, SPIN90-KO results in an increase in total Arp2/3 complex density and phalloidin staining (Fig. 5) (28). These findings are at odds with a simple model where SPIN90 acts as an NPF to localize Arp2/3 complex activation. Adding to this complexity, we found that SPIN90-KO cells are more contractile and yet migrate faster (Fig. 3), which is the opposite of what is typically found (55–57). SPIN90’s SH3 domain localizes to FAs and stunts their maturation when overexpressed (Figs 2I and 4I-K), suggesting an intimate connection between SPIN90 and FAs that could be key to unraveling the role of the Arp2/3 complex in nascent FA assembly (38, 39). Further investigation into SPIN90’s effects on cortical dynamics and structure should help to elucidate the underlying mechanisms.

We found that SPIN90 can enter SF-FA bundles through its DD-ARM domain (Fig. S5), suggesting that Arp2/3 complex activation by this domain results in unbranched filaments that can be bundled into SFs. In contrast, SPIN90 SH3 domain’s enrichment in FAs parallels that of SPIN90 binding partners Nck and palladin (Figs. 2I and S1) (30, 43, 58–61). SPIN90, Nck, palladin, and N-WASP also colocalize in costameres, structures associated with Z-disks analogous to FAs, and are necessary for sarcomere organization in cardiomyocytes (59, 62, 63), highlighting a conserved function across cell and tissue types. Nck proteins bind to phosphorylated FAK in FAs through an SH2 domain and SPIN90’s PR regions through two SH3 domains (30, 59, 64). SPIN90’s SH3 domain binds directly to the PR sequence of palladin, which in turn binds both VASP, an FA actin nucleator, and α-actinin (60, 65, 66). The SPIN90 SH3 likely also binds transiently to PR regions of N-WASP to collaboratively nucleate unbranched actin in FAs (39, 67–69), and it may also bind directly to SFs through the FH1 domain of the formin mDia1 (29, 70). Therefore, there are multiple interactions described in the literature that could explain how each of SPIN90’s domains binds to SF-FA bundles.

In conclusion, while the SPIN90 family’s ability to enhance branched actin assembly is widely acknowledged (24–26, 28), our study raises a new possibility that unbranched SPIN90-Arp2/3 actin filaments are more than just the mothers of branches; they also function within linear SF-FA bundles.

## Materials and Methods

### Tissue culture, genetics, and lentivirus

U2OS (HTB-96, ATCC, RRID: CVCL_0042), 293T (CRL-3216, RRID: CVCL_0063), MEF, CRL-2991, ATCC, RRID: CVCL_U690), NIH-3T3 (CRL-1658, ATCC, RRID: CVCL_0594, and HeLa (RRID: CVCL_0030) were maintained in McCoy’s 5A (catalog #16600082) and DMEM media (#11995065; ThermoFisher, Waltham, MA) at 37°C and 5% CO_2_. Media were supplemented with 10% v/v fetal bovine serum (FBS; ThermoFisher, #A5256701), 1x Antibiotic-Antimycotic (Anti-Anti; ThermoFisher, #15240062), and 1x Glutamax (ThermoFisher, #35050061).

Homozygous amino (N)-terminal CRISPR knock-in U2OS clones for Halo-ArpC1B and Halo-SPIN90 are made by EditCo (Redwood City, CA) using guide RNAs (ArpC1B) CAGGAAGCUGUGGUAGGCCA and (SPIN90) AGCGCGCGGUACAUGAGGCC and confirmed by gene sequencing. Donor sequences encoding HaloTag7 (AQS79244.1) with linkers HRLGKPGLGLQ (ArpC1B) and EPTTEDLYFQSDNAIA (SPIN90) were used for in-frame insertion upstream of A2 of ArpC1B and Y2 of SPIN90 (Fig. S2). Homozygous SPIN90-KO U2OS were made by Ubigene (Guangzhou, P.R. China) using CRISPR guide RNA sequences (G1) GCUGUACGCGUUCCGCUCGGCGG and (G2) AGCGAAGCAGCGCGCACUGGUGG, targeting the first exon of the *NCKIPSD* gene. Each KO clone was confirmed by sequencing to have a specific 71 base pair deletion that causes a frame-shift and early termination resulting in the coding sequence for <u>MYRALYAFR</u>LVAGRAGAQW-stop (retained 9aa SPIN90 sequence underlined, followed by missense residues, Fig. S4).

Lentiviruses were packaged by co-transfection of 27 µg psPX2 and 9 µg PMD.G together with 36 µg expression construct (Table S1) into 293T cells at 40-60% confluency in 56.7 cm^2^ culture dishes using lipofectamine 2000 protocol (Invitrogen, Waltham, MA). After one change of media within 24 h, media supernatants containing viruses were harvested 3 days post-transfection by centrifugation at 100 × g for 5 min and filtration through a 0.45 μm pore-size membrane. Lentivirus aliquots were stored at -80°C. Expression constructs (Table S1) from our study were assembled by GenScript (Piscataway, NJ) while others were obtained from Addgene (Watertown, MA).

### Live cell single molecule tracking experiments

U2OS grown to 40-60% confluency in 9.6 cm^2^ culture chambers were transduced with lentiviruses (40-100 µL) in culture media with 10 µg mL^-1^ polybrene. Alternatively, cells were transfected with 0.5 µg LifeAct-EGFP and 0.3 µg Halo-protein plasmids using lipofectamine 3000 protocol (Invitrogen). After 24 h transduction, cells were re-seeded at 20-30% confluency onto glass bottom, 0.7 cm^2^-per-well imaging slides coated with 10 µg mL^-1^ fibronectin. The next day, cells were stained for 15 min in the 37°C, 5% CO_2_ tissue incubator with 1 nM, 2 nM (for endogenous Halo-ArpC1B), or 20 nM (for endogenous Halo-SPIN90) Janelia Fluor-JFX^650^ Halo ligand (Promega, Fitchburg, WI, #HT1070) in culture media. Cells were then washed twice with culture media and then covered in imaging media [L-15 Leibovitz’s media (ThermoFisher, #11415064) with 10% FBS, 1x Glutamax, 1x Anti-Anti, and 10 mM HEPES pH 7.5]. Next, these samples were mounted into a Nanoimager microscope (ONI, San Diego, CA) that was prewarmed to 37°C and imaged at 5 s frame intervals with 500 ms exposures. The basal cortices of cells were illuminated in TIRF by 488 nm and 640 nm excitation lasers angled to 54.5° at 0.1-0.5 % and 10-16% max power. To capture leading edge dynamics (Fig. S3B), a near-TIRF angle was employed. Micrograph series were typically 500 frames (41:35 m:s) but range from 15 min to 1 h in duration.

Under the above conditions, we selected cells to image and analyze where discrete puncta of approximately uniform size and intensity indicated that single-molecule labeling was achieved. TrackMate was used to localize and track Halo-labeled particles (71). Particles were sub-pixel localized using the Laplacian of Gaussian detector with an estimated object diameter of 0.5 µm and quality thresholds of 15 and above. TrackMate’s simple Linear Assignment Problem tracker was used to assign trajectories, with a max linking distance of 0.3 µm, a gap-closing distance of 0.3 µm, and a max frame gap of 2. Resulting trajectories were filtered with a minimal duration of either 120 s or 300 s to minimize low-specificity associations.

Mean squared displacement (MSD) trajectory analysis was performed using publicly available software (https://github.com/melikelakadamyali/StormAnalysisSoftware) (72). For trajectories with MSD α value ≥ 1.2 (α = slope of log[MSD] / log[t]; time differential, t) that were considered directed, speeds were calculated by the parabolic fit where MSD(t) = a*t + c*t^2^, a = 4*D (diffusion coefficient, D), and c = *v*^2^ (velocity, *v*).

For lifetime statistics (Table 1, Fig. 2E), the cumulative frequency (cf) distributions of trajectory durations (d; cutoff of 12 frames) were fit to double exponentials (*cf*(*d*) = *cf*_0_ + *A_fast_e*^−*k*^*^fastfd^* + *A_slow_e*^−*k*^*^slow^*^d^; A, amplitude; k, rate). Reported lifetimes (τ) were corrected for photobleaching by subtracting the rate of bulk cytoplasmic fluorescence decay of unfused Halo, k_bleach_ (τ_corrected_ = 1 / (k_observed_ – k_bleach_)).

### Wound healing motility

U2OS cells were seeded inside of 2-well inserts on Ibidi 35 mm μ-dishes (Ibidi USA, Fitchburg, WI), and allowed to grow to confluency, before removal of the insert to create the wound gap. The cells were briefly rinsed twice with growth media to remove cell debris. The entire wound gap was imaged at 0, 3, 6, and 10.5 h post-wound on an EVOS M7000 microscope (ThermoFisher) with a 10 × LWD objective under transmitted light. The 10.5 h end time of the wound healing was chosen as regions along the cell fronts in SPIN90 KO cells started to make contact. The images acquired were post-processed using ImageJ. The mean gap distance (*dgap*) was measured by the total area of the gap (tracing the leading cells) divided by the length of the entire gap. The distance traveled by each cell front is the change in half gap distance from the initial (*Δdgap_1/2_* = [*dgap*^t=0h^ – *dgap*^t=n^] / 2). The average cell front speeds were determined by the slopes of linear fits (distance traveled / time). Four independent experiments were conducted, each including a WT and both SPIN90-KO clones.

### Traction force microscopy (TFM) in brief

TFM was performed as described (73). See *Supplemental Materials and Methods* for detail.

### Immunofluorescence (IF)

U2OS were allowed to attach to 10 µg mL^-1^ fibronectin-coated imaging slides for > 2 h before fixation for 15 min at 37°C, 5% CO_2_ in prewarmed PBS (phosphate buffered saline) with 4% w/v paraformaldehyde (Electron Microscopy Sciences, Hatfield, PA). Following three washes with PBS, fixed cells were stored at 4°C. Cells are permeabilized with 0.2% v/v Triton X-100 in blocking buffer (3% w/v BSA in PBS) for 1 h at ambient temperature and then stained with primary antibodies (Table S2) in blocking buffer overnight at 4°C. Cells were washed three times with PBS before incubating with secondary antibodies (Table S2) or 165 nM AlexaFluor^647^ phalloidin (Invitrogen, #A22287) in blocking buffer for 1h at ambient temperature. Before imaging, samples were washed three times with PBS and then covered in imaging buffer (0.56 mg mL^-1^ glucose oxidase, 40 µg mL^-1^ catalase, and 5% w/v glucose in PBS). TIRF, or near-TIRF (Fig. S3A), images were acquired at ambient temperature with 500 ms exposure at fixed 488 and 640 laser powers (0.1-2%). Biological repeat experiments represent individual days when a complete set of WT and SPIN90-KO clones were fixed simultaneously. For each experiment, images were taken in rapid succession (capturing one or more cells) per condition, with the WT divided evenly into two sessions immediately before and after the KO clones.

### Western blot experiments

U2OS were harvested (40 h post-transduction when applicable) and lysed with ice-cold Cell Lysis Buffer (Cell Signaling, Danvers, MA, #9803S) supplemented with 1x cOmplete protease cocktail (Millipore Sigma, Burlington, MA, #04693116001) and 1 mM phenylmethylsulfonyl fluoride (PMSF), clarified by centrifugation at 21,300 × g at 4 °C. Total protein concentrations were measured by Bradford protein assay (ThermoFisher, #1856210). Supernatant fractions were then prepared for SDS-PAGE by the addition of 1x NuPAGE LDS sample buffer (Invitrogen, #NP0007) and 1x NuPAGE Reducing Agent (Invitrogen, #NP0009), and boiling for 7 min at 98 °C.

Lysate samples were then resolved on NuPAGE 4-12% Bis-Tris gels (Invitrogen) and transferred to nitrocellulose membranes. The blots were blocked in Intercept Odyssey TBS (tris buffered saline) blocking buffer (LI-COR, Lincoln, NE, #927-60001) for 1 hr at 25 °C before incubation with primary antibodies (Table S2) at 4 °C overnight in Odyssey TBS blocking buffer with 0.05% Tween-20. Then, the blots were washed three times with TBST buffer (TBS supplemented with 0.05% v/v Tween-20) before incubation with secondary antibodies (Table S2) for 1 hr at 25 °C. The blots were then washed three times with TBST buffer and imaged on LI-COR Odyssey (Fc) Imager.

### Graphing and statistics

GraphPad Prism version 10.6.1 was used for graphing and statistics.

## Supporting information

Mov. S2

## Acknowledgements

We thank Natali Chanaday, Subramanian Ramanathan, and Sophie Travis for helpful comments on the manuscript. We thank the Microscopy Core Research Facility (RRID:SCR_017798) at UNL and Bara Altartouri for microscopy support.

## Author contributions

Writing-original draft: L.W.P and Q.T. Conceptualization: L.W.P., J.R.B, P.W.O, and Q.T. Writing-review and editing: L.W.P. and Q.T. Methodology: L.W.P., J.R.B, P.W.O, and Q.T. Formal analysis: L.W.P., C.L.B., E.L.S., and Q.T. Investigation: L.W.P., C.L.B., E.L.S., A.J.S., and Q.T. Resources: J.R.B, P.W.O, and Q.T. Visualization: L.W.P. and Q.T. Supervision: L.W.P., J.R.B, P.W.O, and Q.T. Data curation: L.W.P., C.L.B., E.L.S., J.R.B, P.W.O, and Q.T. Validation: L.W.P., J.R.B, P.W.O, and Q.T. Funding acquisition: J.R.B, P.W.O, and Q.T. Project administration: L.W.P. and Q.T.

## Data and materials availability

All data needed to evaluate the conclusions in the paper are present in the paper and/or the Supplementary Materials.

## Supplemental Materials and Methods

### Relative basocortical binding of Halo-SPIN90 and subfragments

To evaluate the binding of SPIN90 mutants to the cortical actin network, we measured the number of single molecule tracks that persist at least 5 min as a function of the number of detected spots (the sum of the total localizations). Presumably, under sparse Halo-ligand labeling, the number of spots detected in each cell scales with the target concentration. The result was a linear relationship that reflects sub-saturating binding as a function of concentration, where slope increased with specific cortical binding: full length Halo-SPIN90 (SPIN90^FL^) had a slope of 8.6 ± 0.5 x 10^-4^ tracks spot^-1^ whereas the slope of Halo alone was significantly lower (F-test *p* < 0.0001), 2.5 ± 0.2 x 10^-4^ tracks spot^-1^ (Fig. S5A). Long-lived events of highly-expressed unfused Halo are plausibly due to nonspecific trapping in a dense cortical meshwork or sequestration into organelles (lysosomes, for example). We reasoned that dissecting SPIN90 into fragments of different size and introducing key mutations should resolve whether its cortical binding is specific. Based on the robust distinction between SPIN90^FL^ and Halo alone, we further evaluated this slope parameter in determining relative cortical binding of SPIN90 segments by SMT.

We next acquired the binding slopes of SPIN90 mutants as a fraction of the SPIN90^FL^ condition and correcting for Halo alone (basal cortex binding index, BCBI). The ARM* mutation in SPIN90^FL^, with a BCBI of 0.98, had no effect (*p* = 0.90) on cortical binding (Fig. S5A and B). However, SPIN90^1-110^, containing the SH3 domain and a contiguous region of predicted α-helix, had a BCBI of 1.38, nearly 40% greater than SPIN90^FL^ (*p* = 0.014, Fig. S5A and C). Since SPIN90^1-110^ is 612 aa smaller than SPIN90^FL^, its increased BCBI suggests that size alone does not account for the difference in slopes between SPIN90^FL^ and unfused Halo, hence the majority of long-lived SPIN90^FL^ trajectories occur though specific binding. SPIN90^269-722^ had a BCBI of 0.84 (*p* = 0.13) and moved together with SFs (Fig. S5A and D). When SPIN90^269-722^ had the ARM* mutation or was further truncated to SPIN90^372-722^, which removed the DD domain, the BCBI was reduced to 0.14 and 0.34 (*p* < 0.0001, Fig. S5A). The BCBI of SPIN90^269-722^ being reduced by 0.7 when the charge-neutral ARM* mutation is introduced was further evidence that the SPIN90^269-722^ cortical binding is specific and not an artifact of trapping within the actin mesh.

### Traction force microscopy (TFM) in detail

To prepare the TFM substrate gels, 22×30mm glass coverslips (Electron Microscopy Sciences, #72204-02) were activated in 2% 3-aminopropyltrimethoxysilane (Sigma-Aldrich, #281778) in isopropanol for 10 minutes, washed 4x for 10 minutes in ddH2O, and then treated with 1% glutaraldehyde in ddH_2_O for 30 minutes, washed 3x for 10 minutes in ddH2O, and dried overnight in tinfoil to keep them in a dust-free environment. A polyacrylamide (PAA) gel was prepared using a 5.4 kPa Shear modulus Standard Solution with a final working solution of 7.5% Acrylamide (BioRad, #161-0140) and 0.20% Bis-Acrylamide (Fisher Scientific, #BP1404-250) and polymerized using 10% Ammonium Persulfate (Fisher Scientific, #BP179-25) and TEMED (Fisher Scientific, #BPP150-20). PAA gels additionally contained 0.04 μm fluorescent microspheres (Invitrogen, #F8793). Fibronectin ECM-coupling to the PAA gel was achieved by UV crosslinking of SulfoSANPAH (Fisher Scientific, #PI22589) diluted in DMSO and incubation with 1 mg/mL human fibronectin (Sigma-Aldrich, #FC010) overnight at 4°C. Gels were rinsed 5-10x in sterile PBS.

U2OS cells were transfected with 2 µg total plasmid DNA per 250,000 cells in 100 µL buffer using the Amaxa Nucleofector I electroporator. Transfected cells were then plated on uncoated plastic tissue culture dishes (CELLTREAT, #229661). Twenty-four hours after transfection, cells were lifted, and 5,000 cells were plated on the TFM gel and allowed to adhere overnight. Experiments were performed 48 hours after transfection. Images of cells and fluorescent beads were acquired before cells were removed, then using 0.05% SDS to obtain an unstrained reference bead image.

Traction forces were analyzed using publicly available code (https://github.com/OakesLab/TFM) (1). Prior to image processing, images were flat-field corrected, with the reference bead image aligned to cell-attached bead images. Bead displacement was calculated using an optical flow algorithm in OpenCV (Open-Source Computer Vision Library, https://opencv.org/) with a window size of 8 pixels. Traction stresses were calculated using the Fourier Transform Traction Cytometry approach (2) with a regularization parameter of 3.86 × 10^−6^. The total contractile work, or strain energy, was calculated as one-half the summed product of substrate displacement and traction vectors beneath the cell. Strain energy values were normalized by cell area derived from the Halo channel image.

---

**Fig. S1.**
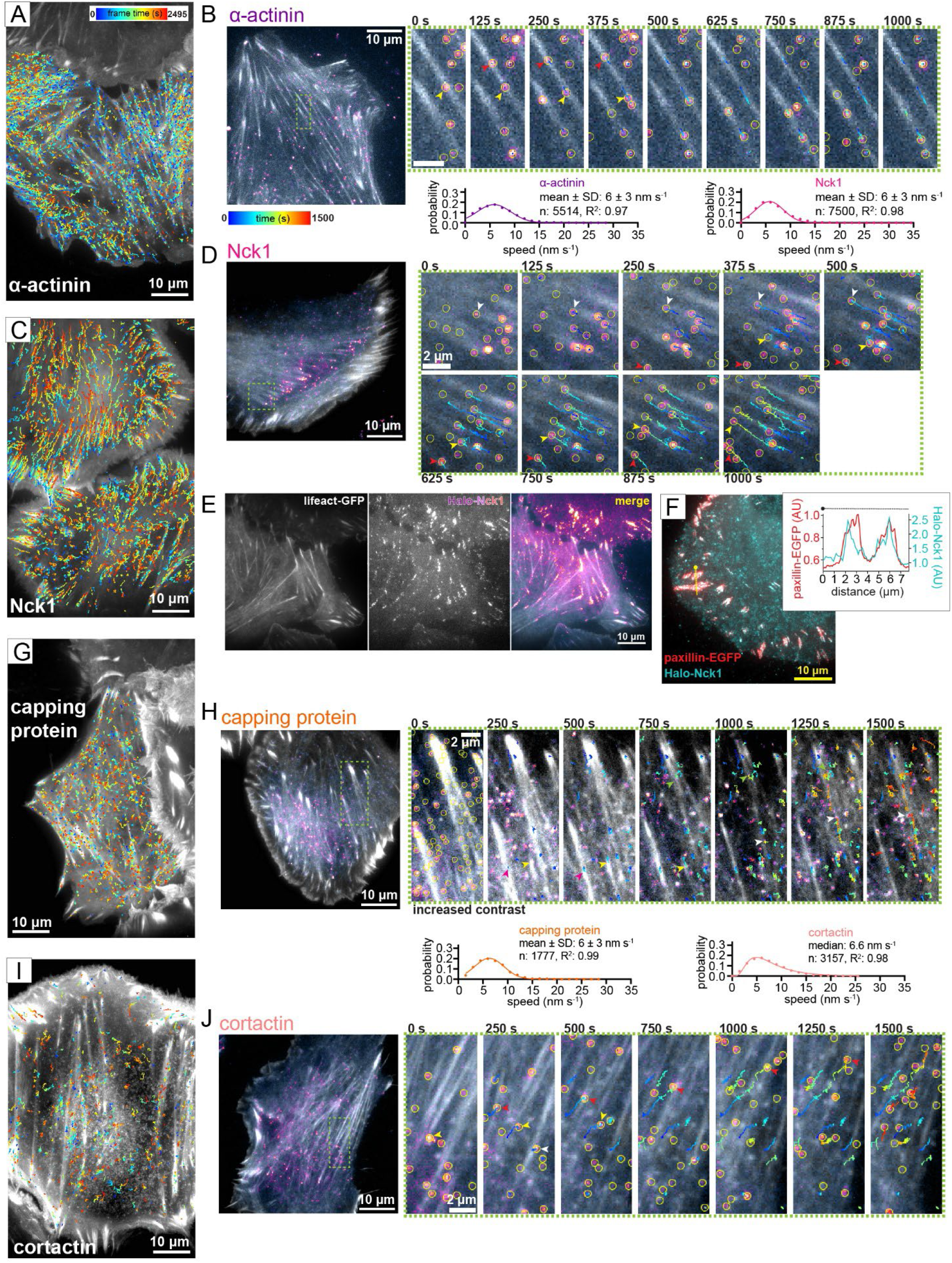
Live cell SMT of α-actinin, Nck1, capping protein, and cortactin. U2OS expressing (**A** and **B**) Halo-α-actinin, (**C-F**) Halo-Nck1, (**G** and **H**) Halo-CAPZ-ϐ or (**I** and **J**) Halo-cortactin were subjected to SMT analysis as in Figs. 1–2. (**A**, **C**, **G**, and **I**) Max. projections of LifeAct-EGFP signal over 500 frames overlaid with trajectories of Halo (unfused) lasting at least 2 min. (**B**, **D**, **H**, and **J**) Example cells (left) and time series (right). Localizations (yellow circles), LifeAct-EGFP (grayscale), and Halo signal (magenta to yellow intensity scale) are shown. For capping protein, localization circles are omitted due to their density obscuring visualization. (**E**) Clusters of Halo-Nck1 in FAs, at the ends of SFs. (**F**) Patches of Halo-Nck1 are associated with paxillin-EGFP shown through maximum projection image (2 min of continuous acquisition at 2 Hz) and line-scan intensity plot. Graphs (**B**, **D**, **H** and **J**) are frequency distributions with N = 4 (α-actinin and Nck1) or 5 (capping protein and cortactin) experiments and indicated numbers of trajectories (n). Solid lines are normal (**B**, **D**, and **H**) and lognormal fits (**J**).

**Fig. S2.**
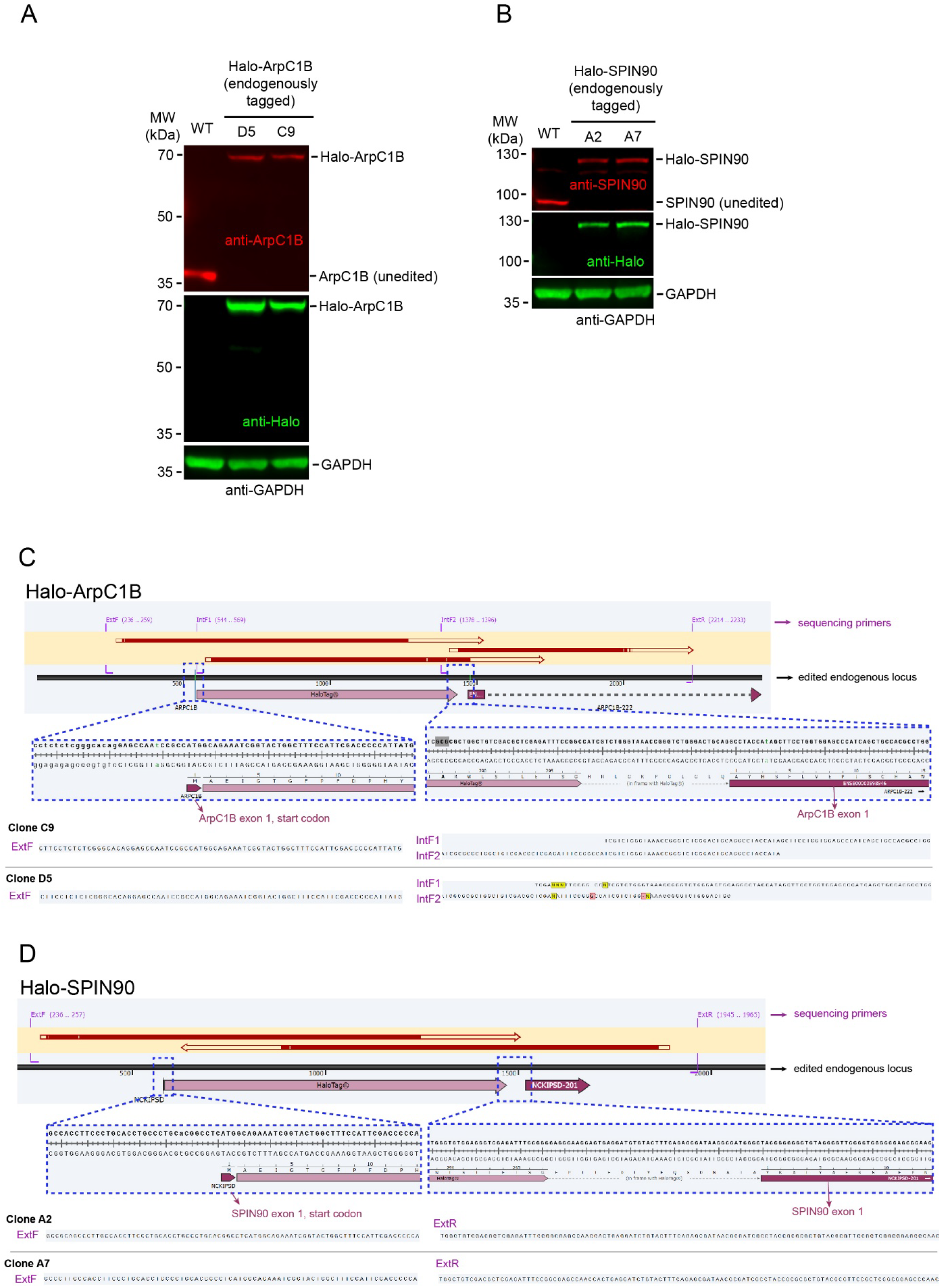
N-terminal Halo-tagging of ArpC1B and SPIN90 at their endogenous loci in U2OS cells. Western blots showing expression of endogenously Halo-tagged (**A**) ArpC1B in two independent clones (D5 and C9), and (**B**) SPIN90 in two independent clones (A2 and A7). The untagged target protein is undetectable in each clone. The cell extract from parental unedited U2OS cells (WT) were recognized by either anti-ArpC1B antibody or SPIN90 antibody, but not by anti-Halo antibody. Each tagged protein shows a ∼33 kDa in molecular weight increase as expected for a Halo-tag. (**C** and **D**) Validation of the integration of Halo-tag N-terminal to the endogenous target loci by Sanger sequencing. Blue dashed boxes zoom in on the sequences at the junctions between the Halo-tag (with an in-frame linker connecting Halo to the target exon 1) and the flanking endogenous sequences. The sequencing data corresponding to the boxed regions are shown for each clone, with the specific primers indicated in purple letters. The annealing positions for the primers and the corresponding sequencing coverage were indicated in the top alignment graphs.

**Fig. S3.**
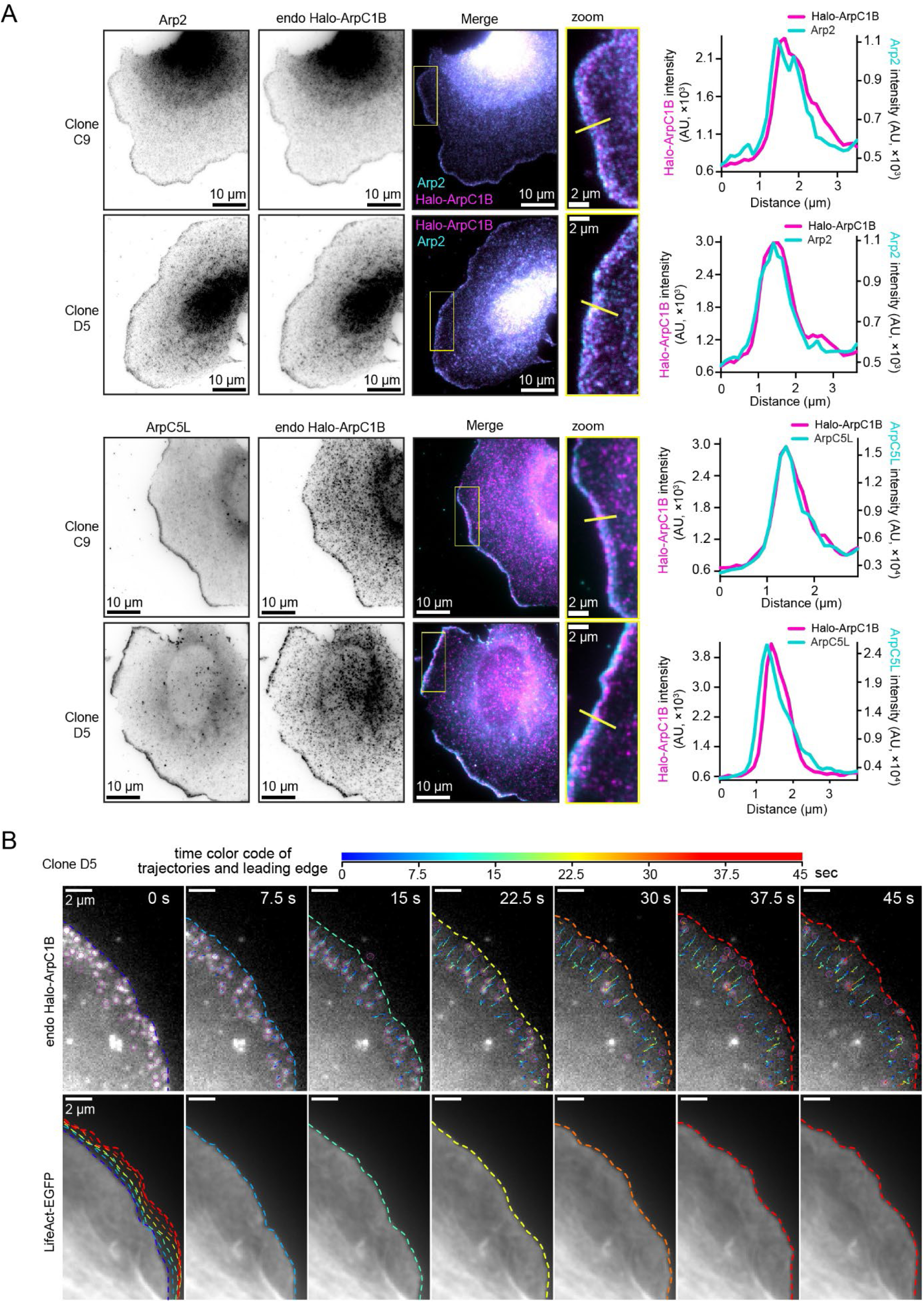
N-terminal Halo-tagging of ArpC1B does not disrupt its localization and dynamics at the leading edge. (**A**) Representative immunofluorescent images of endogenously (endo) tagged Halo-ArpC1B, detected by anti-Halo antibodies, in two independent U2OS clones, C9 and D5, showing colocalization with Arp2 and ArpC5L subunits of Arp2/3 complex at the leading edge. Yellow lines indicate regions selected for linescan intensity in the zoom (yellow rectangles). (**B**) Time series showing single molecules of Halo-ArpC1B (endogenous clone D5) flowing retrograde in the lamellipodium region enriched in LifeAct-EGFP signal over 45 seconds (imaged at 2 Hz resolution). The trajectories and the lamellipodial edge outlines (dashed lines in LifeAct-GFP images) are shown with color-coded time progression. The lower left panel shows all of the outlines to demonstrate the leading edge progression over time.

**Fig. S4.**
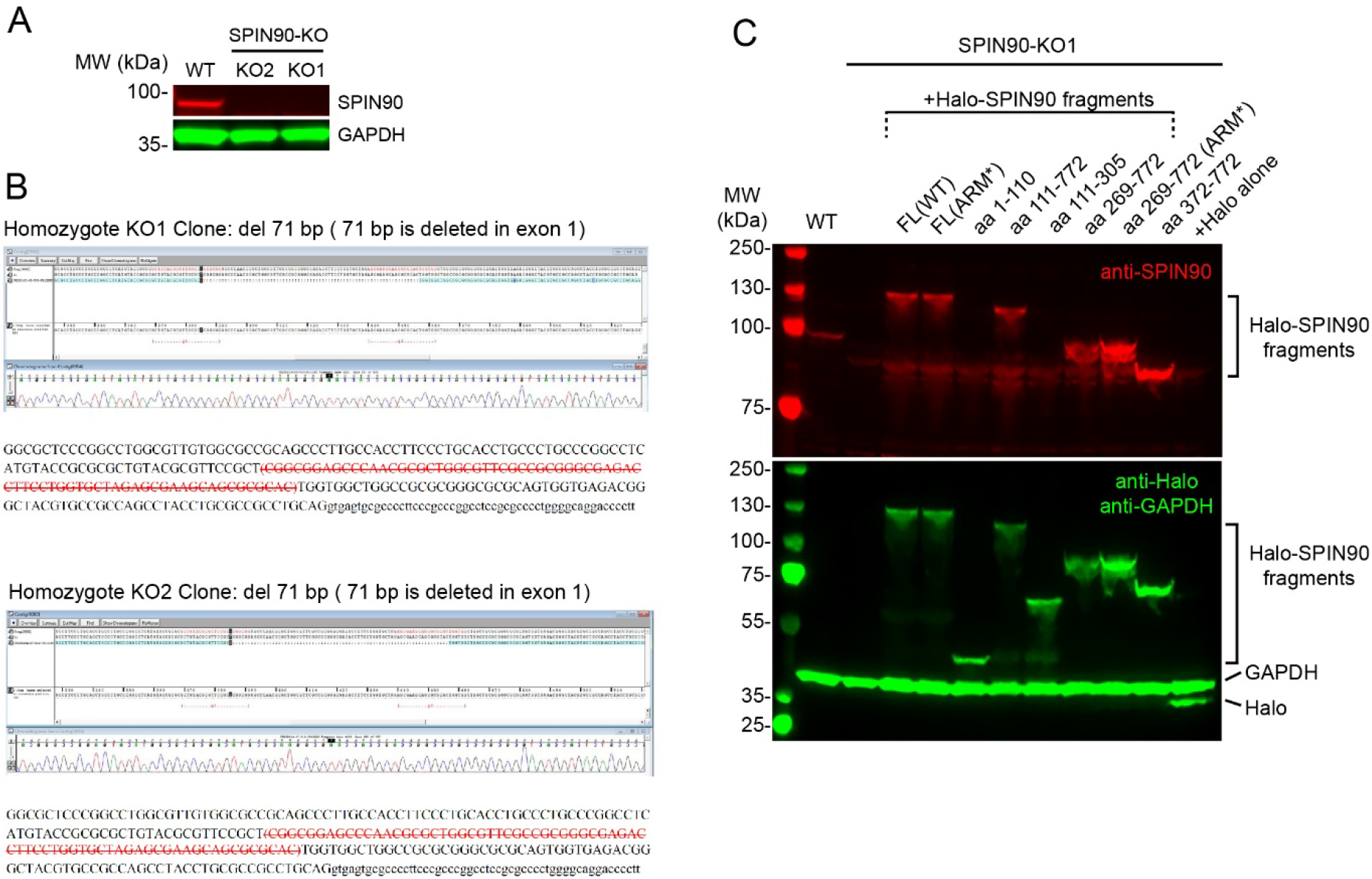
Expression of Halo-SPIN90 constructs in U2OS SPIN90-KO cells. (**A**) Western blot showing lack of SPIN90 expression in two SPIN90-KO cell clones, KO1 and KO2, in comparison to unedited U2OS cells (WT). (**B**) Sanger sequencing showing disrupted endogenous loci of SPIN90 exon 1 regions in KO1 and KO2 clones. The strike-through red uppercase letters are the 71-nt deleted exon sequences. The lowercase letters indicate intron sequence. (**C**) U2OS WT or SPIN90-KO1 cells expressing the indicated SPIN90 construct. The example blot shows various fragments of Halo-SPIN90 resolved at their expected sizes and significant breakdown products are not observed. Certain SPIN90 fragments were not detected by the anti-SPIN90 antibody due to lack of recognizable antigen sequence. Full length (FL) indicates SPIN90^1-722^.

**Fig. S5.**
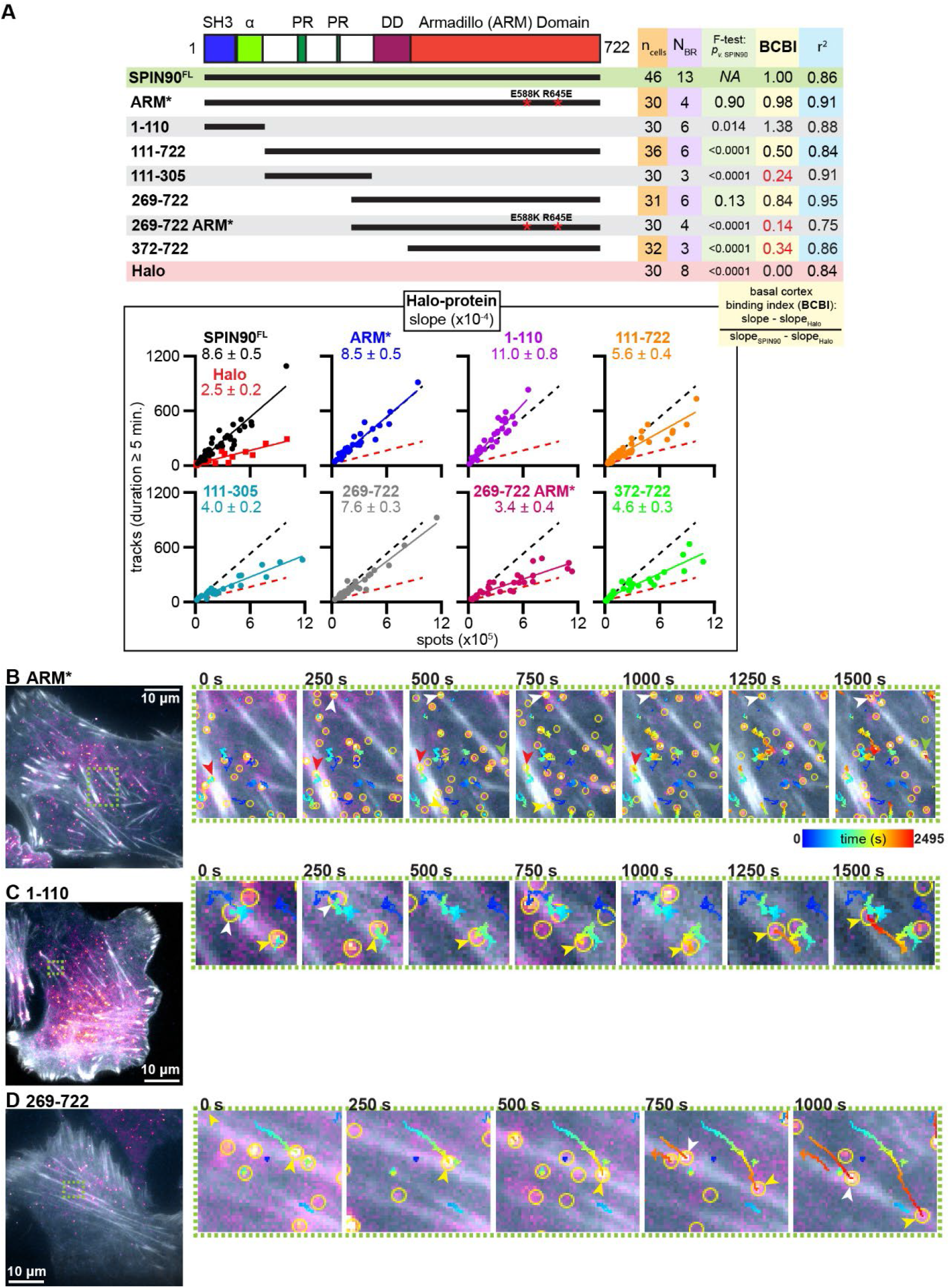
Domain analysis of SPIN90 binding to the basal cortex by SMT. Polypeptides of Halo fused to the indicated amino acid (aa) residues of SPIN90, or Halo alone, were expressed in SPIN90 knockout (KO) U2OS cells and live cell SMT was performed with acquisitions every 5 s for 15 min to 1 h (*Materials and Methods*). (ARM*) ARM domain residues were mutated: E588K and R645E. Variable expression yielded a range in single molecule localizations, or spots, detected by TrackMate (3) within the single molecule labeling regime. See Fig. S4 for Western blot showing expression of all SPIN90 variants. (**A**) The number of trajectories, or tracks, are positively correlated to the number of spots detected. (Above) Data summary and (box below) scatterplots with linear fits (solid lines, R^2^). (BCBI) SPIN90 mutant binding to the basal cortex is shown as a fraction of the slope of full length (FL) SPIN90, correcting for the slope of Halo alone. BCBI below 0.5 are indicated in red letters. Statistics: F-test comparison of slopes with the FL SPIN90. (**B-D**) Example cells (left) and time series (insets, right) showing Halo-labeled molecules and their associated trajectories moving in stress fibers (arrowhead color indicates individual molecules). Representatives, (**B**) 1-722 ARM*, (**C**) 1-110, and (**D**) 269-722, are Halo-SPIN90 fusions that have slopes equal to (*p* > 0.05) or greater than FL SPIN90 (**A**). Localizations (yellow circles), LifeAct-EGFP (grayscale), and Halo signal (magenta to yellow intensity scale) are shown.

**Fig. S6.**
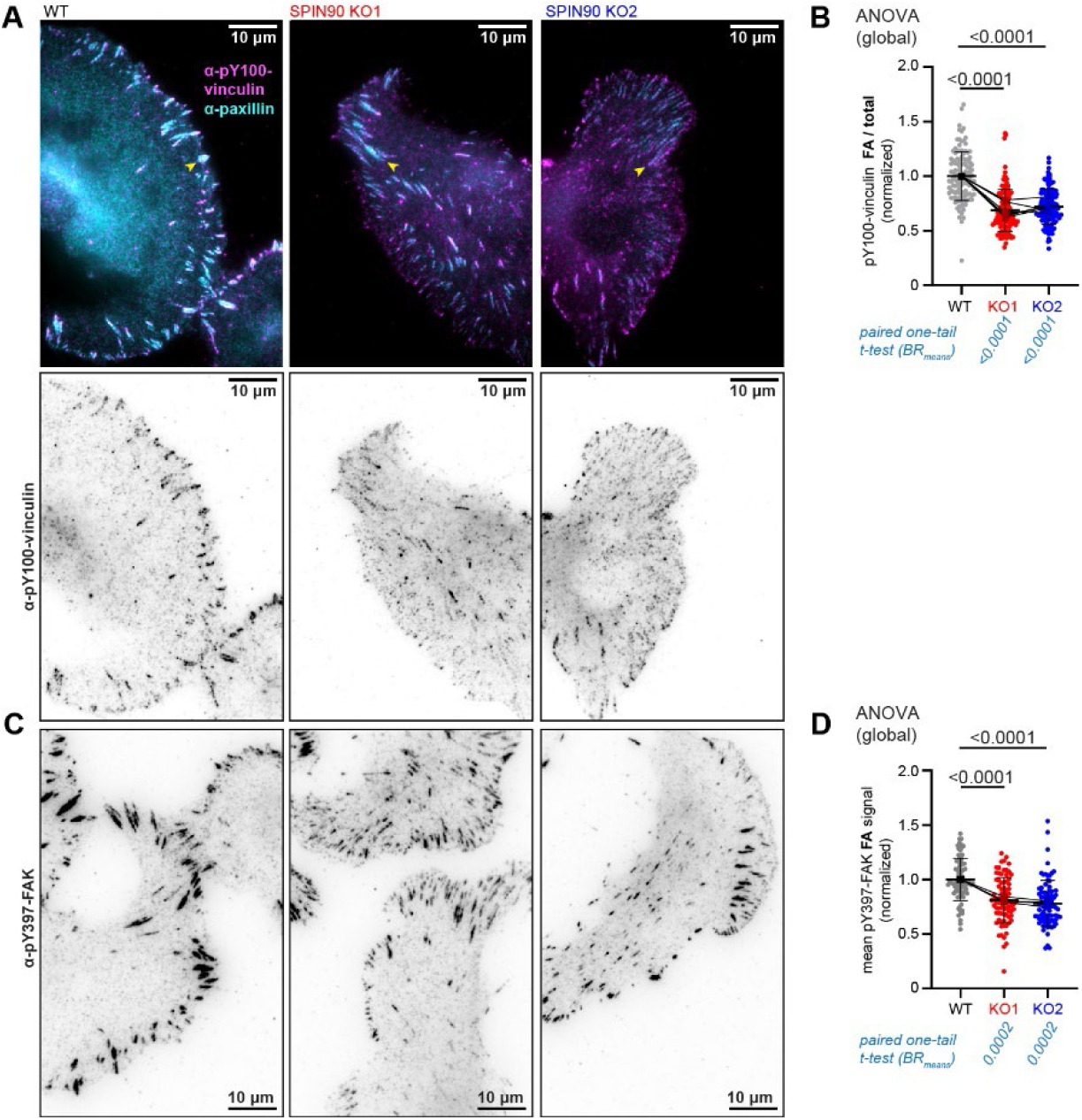
Effects of SPIN90 ablation on phospho-vinculin and -FAK FA staining. WT or SPIN90-KO U2OS subjected to IF analysis using antibodies against phospho-(pY100)-vinculin, paxillin, or phospho-(pY397)-FAK zyxin. (**A**) Example cells imaged in TIRF showing (magenta, Alexa^647^) pY100-vinculin and (cyan, Alexa^488^) paxillin staining. (**B**) The ratio of pY100-vinculin signal in paxillin-stained FAs and total cellular signal (fraction of WT mean). Data are from N_BR_ = 6 experiments with n = 120 images for each condition. (**C**) Example cells stained for pY397-FAK (Alexa^647^). (**D**) Intensity of pY397-FAK. Data are from N_BR_ = 4 experiments and n = 80 images for each condition.

**Table S1.** Lentivirus expression constructs.

| number | tags | linker | protein | residues | ID | species | vector backbone | source |
|---|---|---|---|---|---|---|---|---|
| 102718 | N-DYKDDDDK-Halo | GGGGSGGGSLQ | β-actin | 2-375 | P60709 | human | pUbC | Addgene |
| 84383 | C-EGFP | PPVAT | LifeAct | 1-19 | Q08641 | synthetic | pLenti | Addgene |
| 187688 | C-EGFP | TTLYTKVV | paxillin | 1-559 | AAC38018.1 | chicken | pLenti | Addgene |
| pQT 72 |  |  | Halo | 1-297 | AQS79244.1 | synthetic | pTet-O-FUW | This study |
| pQT 95 | N-Halo | EPTTEDLYFQSDNAIA | nonmuscle myosin regulatory light chain ML12B | 1-172 | O14950 | human | pTet-O-FUW | This study |
| pQT 94 | N-Halo | EPTTEDLYFQSDNAIA | α-actinin ACTN4 | 1-911 | O43707-1 | human | pTet-O-FUW | This study |
| pQT 136 | N-Halo | SGENLYFQGYPYDVPDYASG | SH2/SH3 adapter protein Nck1 | 1-377 | P16333-1 | human | pTet-O-FUW | This study |
| pQT 117 | N-Halo | PAGGSPGLEVNGSGSGSH | capping protein β subunit CAPZB | 1-272 | P47756-2 | human | pTet-O-FUW | This study |
| pQT 81 | N-Halo | HRLGKPGLGLQ | ArpC1B | 1-372 | O15143 | human | pTet-O-FUW | This study |
| pQT 79 | N-Halo | SGLRSRAQASNSAVDGT | cortactin | 1-546 | AAA19689.1 | mouse | pTet-O-FUW | This study |
| pQT 78 | N-Halo | EPTTEDLYFQSDNAIA | SPIN90 | 1-722 | Q9NZQ3-1 | human | pTet-O-FUW | This study |
| pQT 165 | N-HA tag-msfEGFP | EPTTEDLYFQSDNAIA | nonmuscle myosin regulatory light chain ML12B | 1-172 | O14950 | human | pTet-O-FUW | This study |
| pQT 108 | N-Halo | EPTTEDLYFQSDNAIA | SPIN90 FL ARM* E588K R645E | 1-722 | Q9NZQ3-mut | human | pTet-O-FUW | This study |
| pQT 101 | N-Halo | EPTTEDLYFQSDNAIA | SPIN90 1-110 | 1-110 | Q9NZQ3-1 | human | pTet-O-FUW | This study |
| pQT 103 | N-Halo | EPTTEDLYFQSDNAIA | SPIN90 1-305 | 1-305 | Q9NZQ3-1 | human | pTet-O-FUW | This study |
| pQT 104 | N-Halo | EPTTEDLYFQSDNAIA | SPIN90 111-722 | 111-722 | Q9NZQ3-1 | human | pTet-O-FUW | This study |
| pQT 132 | N-Halo | EPTTEDLYFQSDNAIA | SPIN90 111-305 | 111-305 | Q9NZQ3-1 | human | pTet-O-FUW | This study |
| pQT 105 | N-Halo | EPTTEDLYFQSDNAIA | SPIN90 269-722 | 269-722 | Q9NZQ3-1 | human | pTet-O-FUW | This study |
| pQT 130 | N-Halo | EPTTEDLYFQSDNAIA | SPIN90 269-722 ARM* E588K R645E | 269-722 | Q9NZQ3-mut | human | pTet-O-FUW | This study |
| pQT 133 | N-Halo | EPTTEDLYFQSDNAIA | SPIN90 372-722 | 372-722 | Q9NZQ3-1 | human | pTet-O-FUW | This study |

**Table S2.** Antibodies.

| Primary Antibodies |  |  |  |  |  | dilution employed |  |
|---|---|---|---|---|---|---|---|
| **antigen** | **source** | **clonality** | **company** | **catalog number** | **RRID** | **Western blot** | **IF** |
| Arp2 | mouse | mono- | Millipore Sigma, Burlington, MA | A6104 | AB_2221965 | 1:400 | 1:200 |
| Arp3 | mouse | mono- | Millipore Sigma, Burlington, MA | A5979 | AB_476749 | 1:1000 |  |
| ArpC1B | rabbit | poly- | Millipore Sigma, Burlington, MA | HPA004832 | AB_1845044 | 1:500 |  |
| ArpC5L | rabbit | mono- | Abcam, Waltham, MA | AB169763 |  | 1:1000 | 1:250 |
| GAPDH | mouse | mono- | ProteinTech, Rosemont, IL | 60004-1-IG | AB_2107436 | 1:3000 |  |
| HaloTag | mouse | mono- | Promega, Fitchburg, WI | G9211 | AB_2688011 | 1:1000 | 1:250 |
| HaloTag | rabbit | poly- | Promega, Fitchburg, WI | G9281 | AB_713650 |  | 1:250 |
| paxillin | mouse | mono- | BD Biosciences, Franklin Lakes, NJ | 610051 | AB_397463 |  | 1:500 |
| phospho-FAK Y397 | rabbit | mono- | Invitrogen, Carlsbad, CA | 44625G | AB_2533702 |  | 1:250 |
| SPIN90 | rabbit | mono- | Cell Signaling, Danvers, MA | 89661 |  | 1:1000 |  |
| phospho-vinculin Y100 | rabbit | poly- | ThermoFisher, Waltham, MA | 44-1074G | AB_2533566 |  | 1:250 |
| zyxin | mouse | mono- | ThermoFisher, Waltham, MA | MA5-31367 | AB_2787004 |  | 1:500 |
| **Secondary Antibodies** |  |  |  |  |  | **dilution employed** |  |
| **antigen** | **source/clonality** | **conjugate** | **company** | **catalog number** | **RRID** | **Western blot** | **IF** |
| mouse IgG | goat/polyclonal | Alexa Fluor 488 | Invitrogen, Carlsbad, CA | A11001 | AB_2534069 |  | 1:500 |
| mouse IgG | goat/polyclonal | Alexa Fluor 647 | Invitrogen, Carlsbad, CA | A21235 | AB_2535804 |  | 1:500 |
| mouse IgG | goat/polyclonal | IRDye 800CW | LI-COR Biosciences, Lincoln, NE | 926-32210 | AB_621842 | 1:20,000 |  |
| rabbit IgG | goat/polyclonal | Alexa Fluor 488 | Invitrogen, Carlsbad, CA | A11008 | AB_143165 |  | 1:500 |
| rabbit IgG | goat/polyclonal | Alexa Fluor 647 | Invitrogen, Carlsbad, CA | A21244 | AB_2535812 |  | 1:500 |
| rabbit IgG | goat/polyclonal | IRDye 680RD | LI-COR Biosciences, Lincoln, NE | 926-68071 | AB_10956166 | 1:20,000 |  |

**Movie S1.**
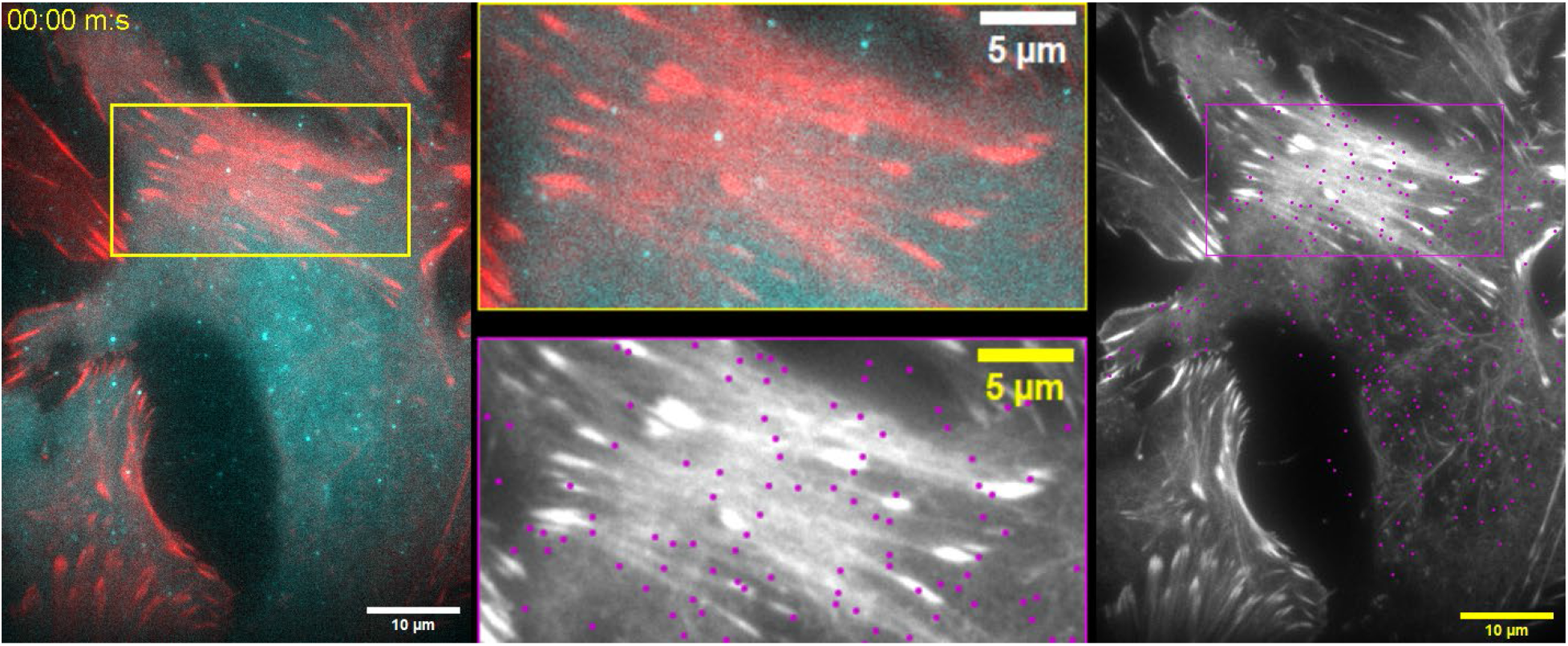
SMT of unfused HaloTag. (Left) TIRF microscopy video showing example cells expressing Halo (cyan) and LifeAct-EGFP (red). (Right) Localizations (magenta dots) and time-coded trajectories overlaid onto the LifeAct-EGFP channel (grayscale) in one of the cells. Zoomed rectangles show a region with dense stress fibers. Trajectory color: blue is 0 s and red is 2,495 s (same as Fig. 1). Playback: 500x speed (capture 0.2 Hz; playback 100 Hz).

**Movie S2.**
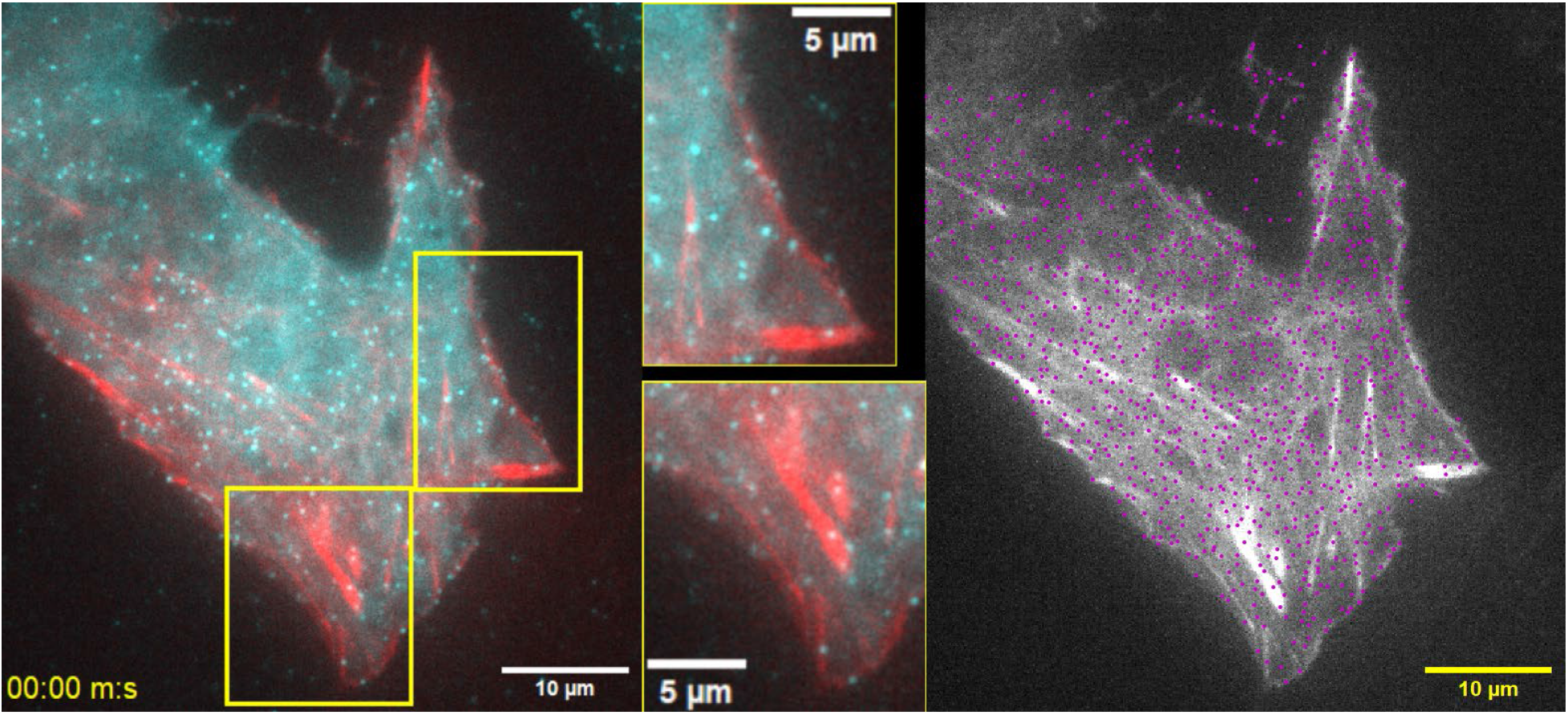
SMT of ectopically expressed Halo-ArpC1B. (Left) TIRF microscopy video showing an example cell expressing Halo-ArpC1B (cyan) and LifeAct-EGFP (red). (Right) Localizations (magenta dots) and time-coded trajectories overlaid onto the LifeAct-EGFP channel (grayscale). Zoomed rectangles show regions were single Halo-ArpC1B molecules are frequently seen flowing together with stress fiber or focal adhesion actin. Trajectory color: blue is 0 s and red is 2,495 s (same as Fig. 1). Playback: 500x speed (capture 0.2 Hz; playback 100 Hz).

**Movie S3.**
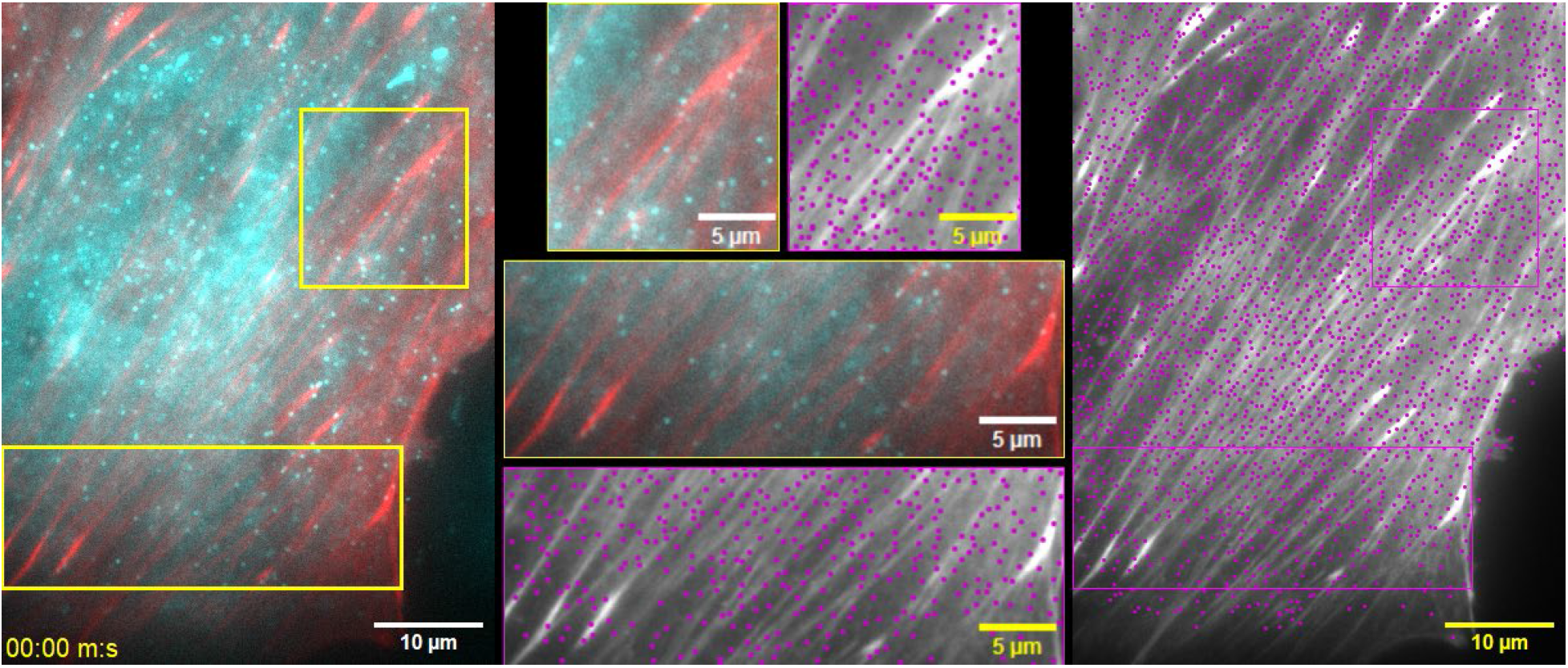
SMT of endogenously Halo-tagged ArpC1B. (Left) TIRF microscopy video showing an example cell with endogenously tagged Halo-ArpC1B (cyan) and expressing LifeAct-EGFP (red). (Right) Localizations (magenta dots) and time-coded trajectories overlaid onto the LifeAct-EGFP channel (grayscale). Zoomed rectangles show regions were single Halo-ArpC1B molecules are frequently seen flowing together with stress fiber or focal adhesion actin. Trajectory color: blue is 0 s and red is 900 s (video duration). Playback: 500x speed (capture 0.2 Hz; playback 100 Hz).

**Movie S4.**
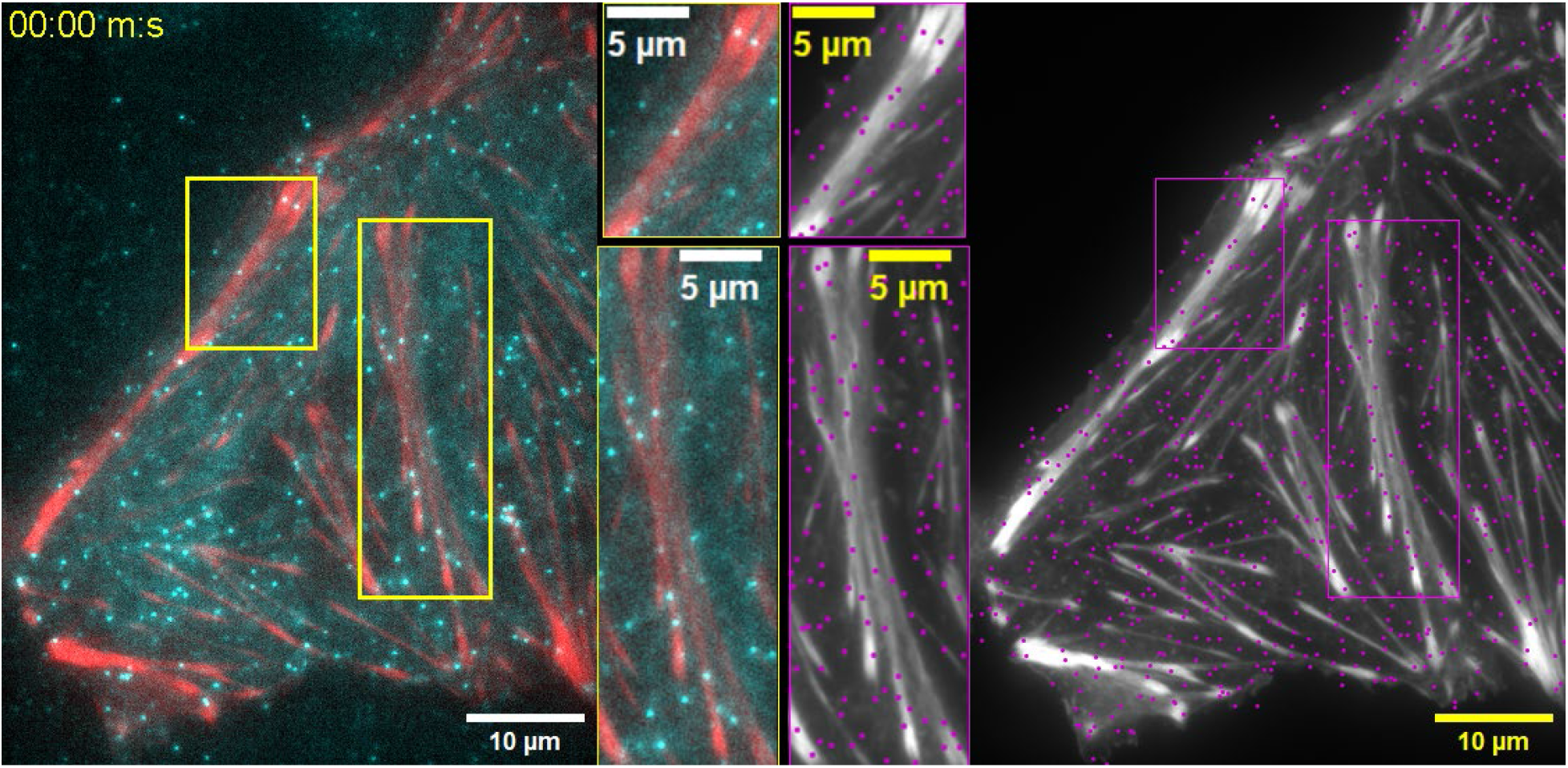
SMT of ectopically expressed Halo-SPIN90. (Left) TIRF microscopy video showing an example cell expressing Halo-SPIN90 (cyan) and LifeAct-EGFP (red). (Right) Localizations (magenta dots) and time-coded trajectories overlaid onto the LifeAct-EGFP channel (grayscale). Zoomed rectangles show regions were single Halo-SPIN90 molecules are frequently seen flowing together with stress fiber or focal adhesion actin. Trajectory color: blue is 0 s and red is 2,495 s (same as Fig. 1). Playback: 500x speed (capture 0.2 Hz; playback 100 Hz).

**Movie S5.**
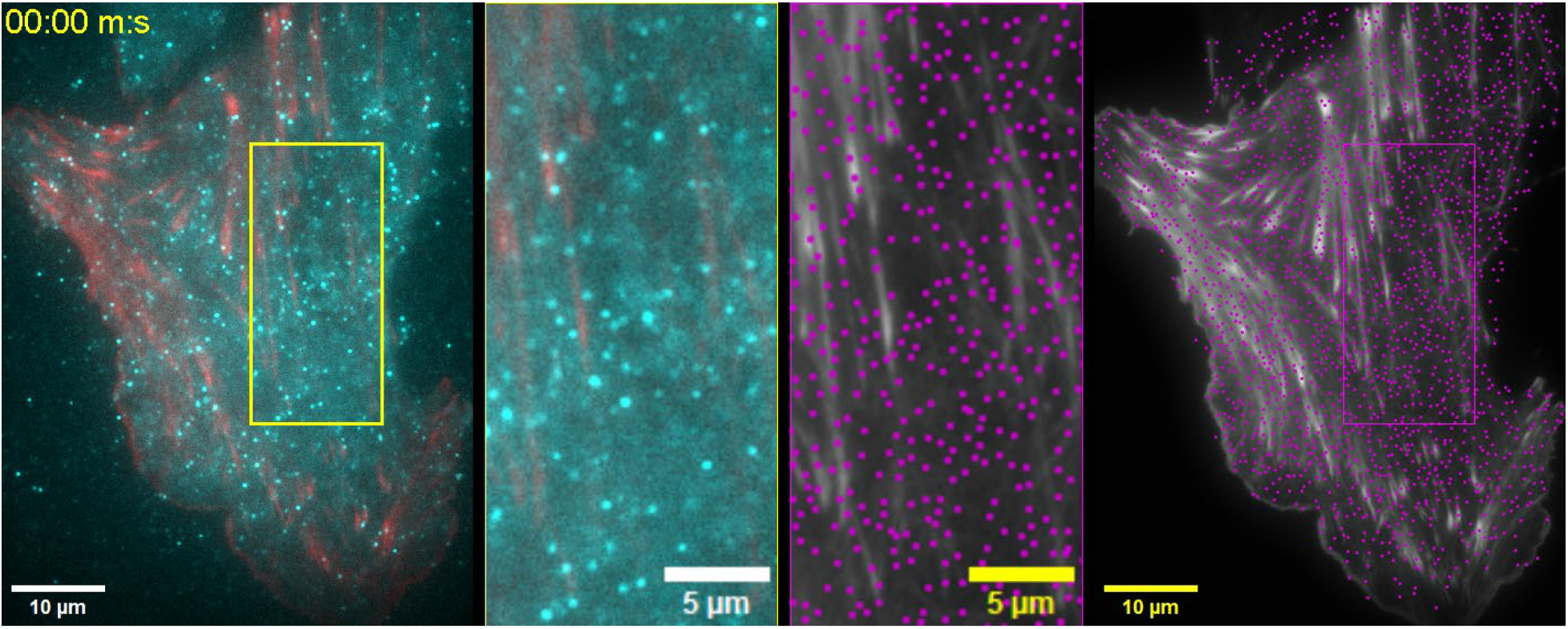
SMT of endogenously Halo-tagged SPIN90. (Left) TIRF microscopy video showing an example cell with endogenously tagged Halo-SPIN90 (cyan) and expressing LifeAct-EGFP (red). (Right) Localizations (magenta dots) and time-coded trajectories overlaid onto the LifeAct-EGFP channel (grayscale). Zoomed rectangles show a region were Halo-SPIN90 molecules are seen moving within stress fibers that contract following assembly. Trajectory color: blue is 0 s and red is 2,400 s (video duration). Playback: 500x speed (capture 0.2 Hz; playback 100 Hz).

**Movie S6.**
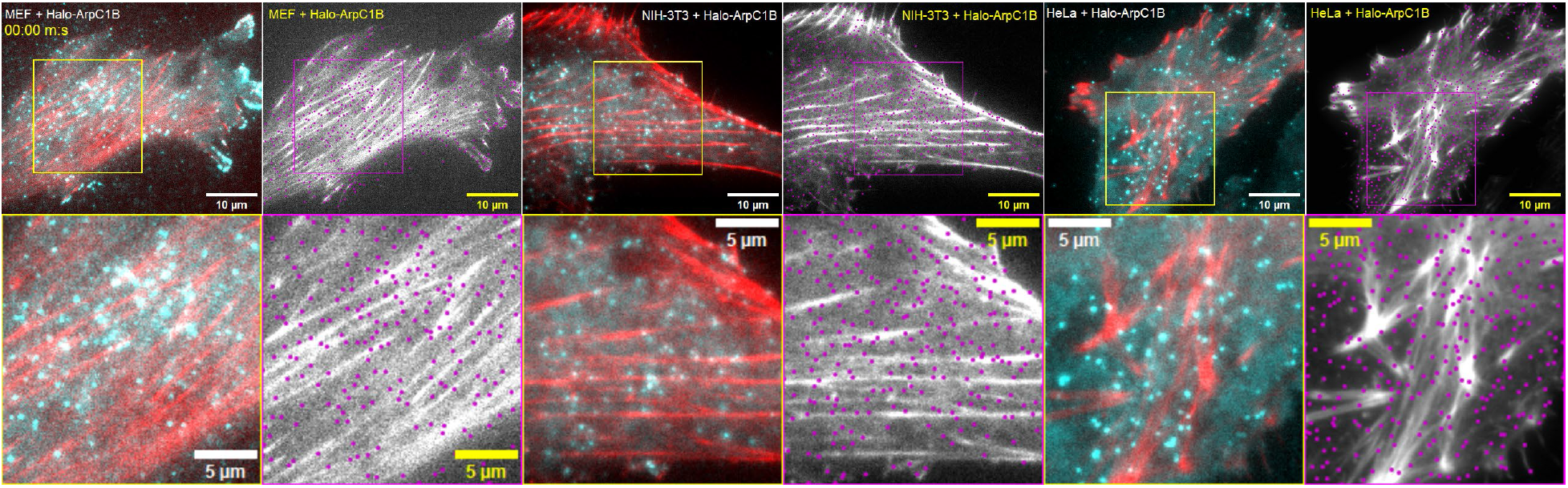
SMT of Halo-ArpC1B ectopically expressed in MEF, NIH-3T3, and HeLa cell lines. (Top) TIRF microscopy videos showing example cells expressing Halo-ArpC1B (cyan) and LifeAct-EGFP (red). Localizations (magenta dots) and time-coded trajectories overlaid onto the LifeAct-EGFP channel (grayscale). Zoomed rectangles (below) show regions were single Halo-ArpC1B molecules are frequently seen flowing together with stress fiber or focal adhesion actin. Trajectory color: blue is 0 s and red is 900 s (video duration). Playback: 125x speed (capture 0.2 Hz; playback 25 Hz).

**Movie S7.**
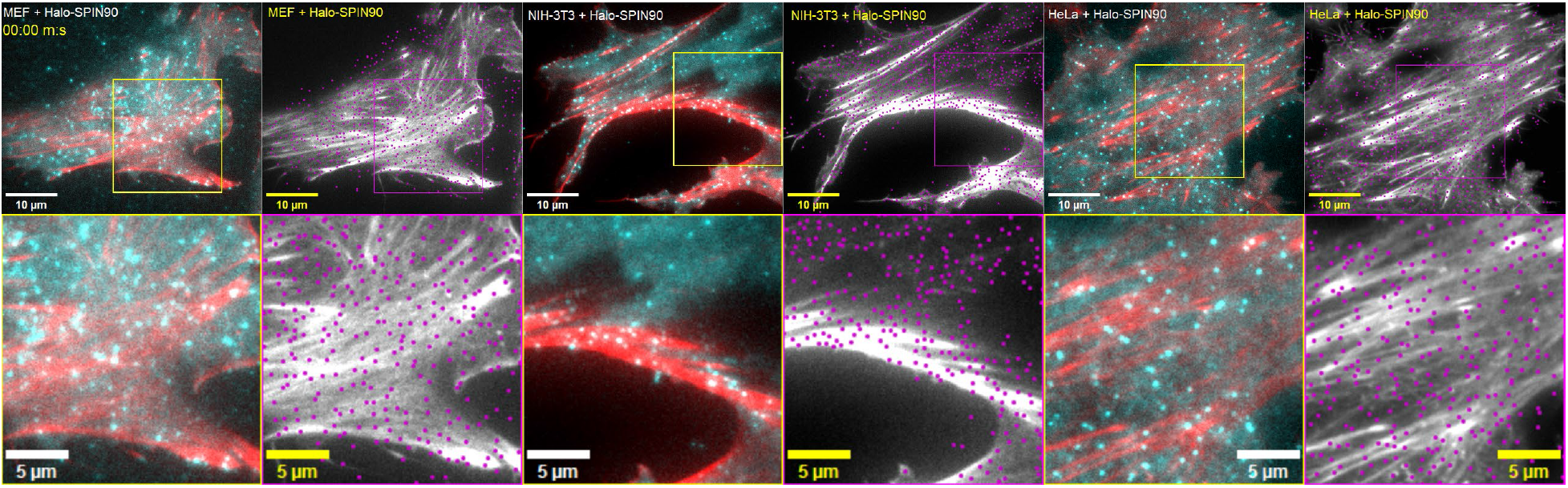
SMT of Halo-SPIN90 ectopically expressed in MEF, NIH-3T3, and HeLa cell lines. (Top) TIRF microscopy videos showing example cells expressing Halo-SPIN90 (cyan) and LifeAct-EGFP (red). Localizations (magenta dots) and time-coded trajectories overlaid onto the LifeAct-EGFP channel (grayscale). Zoomed rectangles (below) show regions were Halo-SPIN90 molecules flow together with stress fiber and focal adhesion actin. Trajectory color: blue is 0 s and red is 900 s (video duration). Playback: 125x speed (capture 0.2 Hz; playback 25 Hz).

## References and Notes

1. L. M. Machesky, S. J. Atkinson, C. Ampe, J. Vandekerckhove, T. D. Pollard, Purification of a cortical complex containing two unconventional actins from Acanthamoeba by affinity chromatography on profilin-agarose. J Cell Biol 127, 107–115 (1994).

2. E. D. Goley, M. D. Welch, The ARP2/3 complex: an actin nucleator comes of age. Nat Rev Mol Cell Biol 7, 713–726 (2006).

3. A. Chaigne, M. H. Verlhac, M. E. Terret, Spindle positioning in mammalian oocytes. Exp Cell Res 318, 1442–1447 (2012).

4. V. Papalazarou, L. M. Machesky, The cell pushes back: The Arp2/3 complex is a key orchestrator of cellular responses to environmental forces. Curr Opin Cell Biol 68, 37–44 (2021).

5. A. M. Gautreau, F. E. Fregoso, G. Simanov, R. Dominguez, Nucleation, stabilization, and disassembly of branched actin networks. Trends Cell Biol 32, 421–432 (2022).

6. A. Szabó, P. Borkúti, Z. Kovács, I. Kristó, P. Vilmos, Recent advances in nuclear actin research. Nucleus 16, 2498643 (2025).

7. I. Rouiller, X. P. Xu, K. J. Amann, C. Egile, S. Nickell, D. Nicastro, R. Li, T. D. Pollard, N. Volkmann, D. Hanein, The structural basis of actin filament branching by the Arp2/3 complex. J Cell Biol 180, 887–895 (2008).

8. M. Shaaban, S. Chowdhury, B. J. Nolen, Cryo-EM reveals the transition of Arp2/3 complex from inactive to nucleation-competent state. Nat Struct Mol Biol 27, 1009–1016 (2020).

9. F. Fäßler, G. Dimchev, V. V. Hodirnau, W. Wan, F. K. M. Schur, Cryo-electron tomography structure of Arp2/3 complex in cells reveals new insights into the branch junction. Nat Commun 11, 6437 (2020).

10. B. Ding, H. Y. Narvaez-Ortiz, Y. Singh, G. M. Hocky, S. Chowdhury, B. J. Nolen, Structure of Arp2/3 complex at a branched actin filament junction resolved by single-particle cryo-electron microscopy. Proc Natl Acad Sci U S A 119, e2202723119 (2022).

11. S. Z. Chou, M. Chatterjee, T. D. Pollard, Mechanism of actin filament branch formation by Arp2/3 complex revealed by a high-resolution cryo-EM structureof the branch junction. Proc Natl Acad Sci U S A 119, e2206722119 (2022).

12. T. E. B. Stradal, M. Boiero Sanders, P. Bieling, Arp2/3-complex regulation - Novel insights and open questions. Curr Opin Cell Biol 95, 102565 (2025).

13. S. B. Padrick, L. K. Doolittle, C. A. Brautigam, D. S. King, M. K. Rosen, Arp2/3 complex is bound and activated by two WASP proteins. Proc Natl Acad Sci U S A 108, E472–479 (2011).

14. M. Rodnick-Smith, Q. Luan, S. L. Liu, B. J. Nolen, Role and structural mechanism of WASP-triggered conformational changes in branched actin filament nucleation by Arp2/3 complex. Proc Natl Acad Sci U S A 113, E3834–3843 (2016).

15. Q. Luan, A. Zelter, M. J. MacCoss, T. N. Davis, B. J. Nolen, Identification of Wiskott-Aldrich syndrome protein (WASP) binding sites on the branched actin filament nucleator Arp2/3 complex. Proc Natl Acad Sci U S A 115, E1409–e1418 (2018).

16. A. Zimmet, T. Van Eeuwen, M. Boczkowska, G. Rebowski, K. Murakami, R. Dominguez, Cryo-EM structure of NPF-bound human Arp2/3 complex and activation mechanism. Sci Adv 6 (2020).

17. A. J. Saks, K. R. Barrie, G. Rebowski, R. Dominguez, NPF binding to Arp2 is allosterically linked to the release of ArpC5’s N-terminal tail and conformational changes in Arp2/3 complex. Proc Natl Acad Sci U S A 122, e2421557122 (2025).

18. A. R. Wagner, Q. Luan, S. L. Liu, B. J. Nolen, Dip1 defines a class of Arp2/3 complex activators that function without preformed actin filaments. Curr Biol 23, 1990–1998 (2013).

19. Q. Luan, S. L. Liu, L. A. Helgeson, B. J. Nolen, Structure of the nucleation-promoting factor SPIN90 bound to the actin filament nucleator Arp2/3 complex. Embo j 37 (2018).

20. C. J. Balzer, A. R. Wagner, L. A. Helgeson, B. J. Nolen, Dip1 Co-opts Features of Branching Nucleation to Create Linear Actin Filaments that Activate WASP-Bound Arp2/3 Complex. Curr Biol 28, 3886–3891.e3884 (2018).

21. T. Liu, L. Cao, M. Mladenov, G. Romet-Lemonne, M. Way, C. A. Moores, Arp2/3-mediated bidirectional actin assembly by SPIN90 dimers. Nat Struct Mol Biol 32, 2262–2271 (2025).

22. J. Francis, A. K. Pathri, K. T. Shyam, S. Sripada, R. Mitra, H. Y. Narvaez-Ortiz, K. V. Eliyan, B. J. Nolen, S. Chowdhury, Activation of Arp2/3 complex by a SPIN90 dimer in linear actin-filament nucleation. Nat Struct Mol Biol 32, 2272–2284 (2025).

23. L. Cao, F. Ghasemi, M. Way, A. Jégou, G. Romet-Lemonne, Regulation of branched versus linear Arp2/3-generated actin filaments. Embo j 42, e113008 (2023).

24. R. Basu, F. Chang, Characterization of dip1p reveals a switch in Arp2/3-dependent actin assembly for fission yeast endocytosis. Curr Biol 21, 905–916 (2011).

25. C. J. Balzer, M. L. James, H. Y. Narvaez-Ortiz, L. A. Helgeson, V. Sirotkin, B. J. Nolen, Synergy between Wsp1 and Dip1 may initiate assembly of endocytic actin networks. Elife 9 (2020).

26. D. J. Kim, S. H. Kim, C. S. Lim, K. Y. Choi, C. S. Park, B. H. Sung, M. G. Yeo, S. Chang, J. K. Kim, W. K. Song, Interaction of SPIN90 with the Arp2/3 complex mediates lamellipodia and actin comet tail formation. J Biol Chem 281, 617–625 (2006).

27. T. Fukumi-Tominaga, Y. Mori, A. Matsuura, K. Kaneko, M. Matsui, M. Ogata, M. Tominaga, DIP/WISH-deficient mice reveal Dia- and N-WASP-interacting protein as a regulator of cytoskeletal dynamics in embryonic fibroblasts. Genes Cells 14, 1197–1207 (2009).

28. L. Cao, A. Basant, M. Mladenov, N. Kogata, A. Jegou, G. Romet-Lemonne, S. Brasselet, M. Mavrakis, M. Way, SPIN90 modulates the architecture of lamellipodial actin in an ARPC5L dependent fashion. bioRxiv 10.64898/2025.12.01.691495, 2025.2012.2001.691495 (2026).

29. L. Cao, A. Yonis, M. Vaghela, E. H. Barriga, P. Chugh, M. B. Smith, J. Maufront, G. Lavoie, A. Méant, E. Ferber, M. Bovellan, A. Alberts, A. Bertin, R. Mayor, E. K. Paluch, P. P. Roux, A. Jégou, G. Romet-Lemonne, G. Charras, SPIN90 associates with mDia1 and the Arp2/3 complex to regulate cortical actin organization. Nat Cell Biol 22, 803–814 (2020).

30. C. S. Lim, S. H. Kim, J. G. Jung, J. K. Kim, W. K. Song, Regulation of SPIN90 phosphorylation and interaction with Nck by ERK and cell adhesion. J Biol Chem 278, 52116–52123 (2003).

31. M. Bailly, F. Macaluso, M. Cammer, A. Chan, J. E. Segall, J. S. Condeelis, Relationship between Arp2/3 complex and the barbed ends of actin filaments at the leading edge of carcinoma cells after epidermal growth factor stimulation. J Cell Biol 145, 331–345 (1999).

32. T. M. Svitkina, Actin Cell Cortex: Structure and Molecular Organization. Trends Cell Biol 30, 556–565 (2020).

33. L. B. Case, C. M. Waterman, Integration of actin dynamics and cell adhesion by a three-dimensional, mechanosensitive molecular clutch. Nat Cell Biol 17, 955–963 (2015).

34. N. Billington, A. Wang, J. Mao, R. S. Adelstein, J. R. Sellers, Characterization of three full-length human nonmuscle myosin II paralogs. J Biol Chem 288, 33398–33410 (2013).

35. M. A. Quintanilla, J. A. Hammer, J. R. Beach, Non-muscle myosin 2 at a glance. J Cell Sci 136 (2023).

36. P. Hotulainen, P. Lappalainen, Stress fibers are generated by two distinct actin assembly mechanisms in motile cells. J Cell Biol 173, 383–394 (2006).

37. V. Dimchev, I. Lahmann, S. A. Koestler, F. Kage, G. Dimchev, A. Steffen, T. E. B. Stradal, F. Vauti, H. H. Arnold, K. Rottner, Induced Arp2/3 Complex Depletion Increases FMNL2/3 Formin Expression and Filopodia Formation. Front Cell Dev Biol 9, 634708 (2021).

38. Y. Beckham, R. J. Vasquez, J. Stricker, K. Sayegh, C. Campillo, M. L. Gardel, Arp2/3 inhibition induces amoeboid-like protrusions in MCF10A epithelial cells by reduced cytoskeletal-membrane coupling and focal adhesion assembly. PLoS One 9, e100943 (2014).

39. V. Swaminathan, R. S. Fischer, C. M. Waterman, The FAK-Arp2/3 interaction promotes leading edge advance and haptosensing by coupling nascent adhesions to lamellipodia actin. Mol Biol Cell 27, 1085–1100 (2016).

40. J. R. Beach, K. S. Bruun, L. Shao, D. Li, Z. Swider, K. Remmert, Y. Zhang, M. A. Conti, R. S. Adelstein, N. M. Rusan, E. Betzig, J. A. Hammer, Actin dynamics and competition for myosin monomer govern the sequential amplification of myosin filaments. Nat Cell Biol 19, 85–93 (2017).

41. J. C. M. Meiring, N. S. Bryce, Y. Wang, M. H. Taft, D. J. Manstein, S. Liu Lau, J. Stear, E. C. Hardeman, P. W. Gunning, Co-polymers of Actin and Tropomyosin Account for a Major Fraction of the Human Actin Cytoskeleton. Curr Biol 28, 2331–2337.e2335 (2018).

42. A. M. Fenix, A. C. Neininger, N. Taneja, K. Hyde, M. R. Visetsouk, R. J. Garde, B. Liu, B. R. Nixon, A. E. Manalo, J. R. Becker, S. W. Crawley, D. M. Bader, M. J. Tyska, Q. Liu, J. H. Gutzman, D. T. Burnette, Muscle-specific stress fibers give rise to sarcomeres in cardiomyocytes. Elife 7 (2018).

43. S. P. Chaki, R. Barhoumi, M. E. Berginski, H. Sreenivasappa, A. Trache, S. M. Gomez, G. M. Rivera, Nck enables directional cell migration through the coordination of polarized membrane protrusion with adhesion dynamics. J Cell Sci 126, 1637–1649 (2013).

44. M. L. Gardel, B. Sabass, L. Ji, G. Danuser, U. S. Schwarz, C. M. Waterman, Traction stress in focal adhesions correlates biphasically with actin retrograde flow speed. J Cell Biol 183, 999–1005 (2008).

45. G. Danuser, C. M. Waterman-Storer, Quantitative fluorescent speckle microscopy of cytoskeleton dynamics. Annu Rev Biophys Biomol Struct 35, 361–387 (2006).

46. F. P. Lai, M. Szczodrak, J. Block, J. Faix, D. Breitsprecher, H. G. Mannherz, T. E. Stradal, G. A. Dunn, J. V. Small, K. Rottner, Arp2/3 complex interactions and actin network turnover in lamellipodia. Embo j 27, 982–992 (2008).

47. M. A. Wear, J. A. Cooper, Capping protein: new insights into mechanism and regulation. Trends Biochem Sci 29, 418–428 (2004).

48. T. Miyoshi, T. Tsuji, C. Higashida, M. Hertzog, A. Fujita, S. Narumiya, G. Scita, N. Watanabe, Actin turnover-dependent fast dissociation of capping protein in the dendritic nucleation actin network: evidence of frequent filament severing. J Cell Biol 175, 947–955 (2006).

49. M. Schnoor, T. E. Stradal, K. Rottner, Cortactin: Cell Functions of A Multifaceted Actin-Binding Protein. Trends Cell Biol 28, 79–98 (2018).

50. T. Liu, L. Cao, M. Mladenov, A. Jegou, M. Way, C. A. Moores, Cortactin stabilizes actin branches by bridging activated Arp2/3 to its nucleated actin filament. Nat Struct Mol Biol 31, 801–809 (2024).

51. J. V. Abella, C. Galloni, J. Pernier, D. J. Barry, S. Kjær, M. F. Carlier, M. Way, Isoform diversity in the Arp2/3 complex determines actin filament dynamics. Nat Cell Biol 18, 76–86 (2016).

52. M. Xu, D. M. Rutkowski, G. Rebowski, M. Boczkowska, L. W. Pollard, R. Dominguez, D. Vavylonis, E. M. Ostap, Myosin-I synergizes with Arp2/3 complex to enhance the pushing forces of branched actin networks. Sci Adv 10, eado5788 (2024).

53. A. V. Hill, The heat of shortening and the dynamic constants of muscle. Proceedings of the Royal Society of London. B. Biological Sciences 126, 136–195 (1938).

54. P. Bleicher, D. Han, N. Billington, R. Hart, C. A. Combs, S. Qazi, I. Chandrasekar, J. R. Knutson, J. R. Sellers, Effect of Load on Non-Muscle Myosin 2 Paralog Filaments in a Biomimetic Contractile Actin Array. Small 22, e07772 (2026).

55. Z. Liu, L. A. van Grunsven, E. Van Rossen, B. Schroyen, J. P. Timmermans, A. Geerts, H. Reynaert, Blebbistatin inhibits contraction and accelerates migration in mouse hepatic stellate cells. Br J Pharmacol 159, 304–315 (2010).

56. A. Leal-Egaña, G. Letort, J. L. Martiel, A. Christ, T. Vignaud, C. Roelants, O. Filhol, M. Théry, The size-speed-force relationship governs migratory cell response to tumorigenic factors. Mol Biol Cell 28, 1612–1621 (2017).

57. D. E. Mason, J. M. Collins, J. H. Dawahare, T. D. Nguyen, Y. Lin, S. L. Voytik-Harbin, P. Zorlutuna, M. C. Yoder, J. D. Boerckel, YAP and TAZ limit cytoskeletal and focal adhesion maturation to enable persistent cell motility. J Cell Biol 218, 1369–1389 (2019).

58. M. M. Parast, C. A. Otey, Characterization of Palladin, a Novel Protein Localized to Stress Fibers and Cell Adhesions. Journal of Cell Biology 150, 643–656 (2000).

59. C. S. Lim, E. S. Park, D. J. Kim, Y. H. Song, S. H. Eom, J. S. Chun, J. H. Kim, J. K. Kim, D. Park, W. K. Song, SPIN90 (SH3 protein interacting with Nck, 90 kDa), an adaptor protein that is developmentally regulated during cardiac myocyte differentiation. J Biol Chem 276, 12871–12878 (2001).

60. M. Rönty, A. Taivainen, L. Heiska, C. Otey, E. Ehler, W. K. Song, O. Carpen, Palladin interacts with SH3 domains of SPIN90 and Src and is required for Src-induced cytoskeletal remodeling. Exp Cell Res 313, 2575–2585 (2007).

61. M. Azatov, S. M. Goicoechea, C. A. Otey, A. Upadhyaya, The actin crosslinking protein palladin modulates force generation and mechanosensitivity of tumor associated fibroblasts. Sci Rep 6, 28805 (2016).

62. K. Takano, H. Watanabe-Takano, S. Suetsugu, S. Kurita, K. Tsujita, S. Kimura, T. Karatsu, T. Takenawa, T. Endo, Nebulin and N-WASP cooperate to cause IGF-1-induced sarcomeric actin filament formation. Science 330, 1536–1540 (2010).

63. G. Mastrototaro, P. Carullo, J. Zhang, B. Scellini, N. Piroddi, S. Nemska, M. C. Filomena, S. Serio, C. A. Otey, C. Tesi, F. Emrich, W. A. Linke, C. Poggesi, S. Boncompagni, M.-L. Bang, Ablation of palladin in adult heart causes dilated cardiomyopathy associated with intercalated disc abnormalities. eLife 12, e78629 (2023).

64. M. Alfaidi, M. L. Scott, A. W. Orr, Sinner or Saint?: Nck Adaptor Proteins in Vascular Biology. Front Cell Dev Biol 9, 688388 (2021).

65. M. Boukhelifa, M. M. Parast, J. E. Bear, F. B. Gertler, C. A. Otey, Palladin is a novel binding partner for Ena/VASP family members. Cell Motility 58, 17–29 (2004).

66. M. Rönty, A. Taivainen, M. Moza, C. A. Otey, O. Carpén, Molecular analysis of the interaction between palladin and alpha-actinin. FEBS Lett 566, 30–34 (2004).

67. M. Fukuoka, S. Suetsugu, H. Miki, K. Fukami, T. Endo, T. Takenawa, A novel neural Wiskott-Aldrich syndrome protein (N-WASP) binding protein, WISH, induces Arp2/3 complex activation independent of Cdc42. J Cell Biol 152, 471–482 (2001).

68. X. Wu, S. Suetsugu, L. A. Cooper, T. Takenawa, J. L. Guan, Focal adhesion kinase regulation of N-WASP subcellular localization and function. J Biol Chem 279, 9565–9576 (2004).

69. A. Misra, R. P. Lim, Z. Wu, T. Thanabalu, N-WASP plays a critical role in fibroblast adhesion and spreading. Biochem Biophys Res Commun 364, 908–912 (2007).

70. K. M. Eisenmann, E. S. Harris, S. M. Kitchen, H. A. Holman, H. N. Higgs, A. S. Alberts, Dia-interacting protein modulates formin-mediated actin assembly at the cell cortex. Curr Biol 17, 579–591 (2007).

71. J. Y. Tinevez, N. Perry, J. Schindelin, G. M. Hoopes, G. D. Reynolds, E. Laplantine, S. Y. Bednarek, S. L. Shorte, K. W. Eliceiri, TrackMate: An open and extensible platform for single-particle tracking. Methods 115, 80–90 (2017).

72. Q. Tang, S. Sensale, C. Bond, J. Xing, A. Qiao, S. Hugelier, A. Arab, G. Arya, M. Lakadamyali, Interplay between stochastic enzyme activity and microtubule stability drives detyrosination enrichment on microtubule subsets. Curr Biol 33, 5169–5184.e5168 (2023).

73. M. S. Schmitt, J. Colen, S. Sala, J. Devany, S. Seetharaman, A. Caillier, M. L. Gardel, P. W. Oakes, V. Vitelli, Machine learning interpretable models of cell mechanics from protein images. Cell 187, 481–494.e424 (2024).

## Supplemental References

1. Schmitt, M. S., Colen, J., Sala, S., Devany, J., Seetharaman, S., Caillier, A. et al. (2024) Machine learning interpretable models of cell mechanics from protein images Cell 187, 481–494.e424

2. Huang, Y., Schell, C., Huber, T. B., Şimşek, A. N., Hersch, N., Merkel, R., et al. (2019) Traction force microscopy with optimized regularization and automated Bayesian parameter selection for comparing cells Sci Rep 9, 539

3. Tinevez, J. Y., Perry, N., Schindelin, J., Hoopes, G. M., Reynolds, G. D., Laplantine, E. et al. (2017) TrackMate: An open and extensible platform for single-particle tracking Methods 115, 80–90

